# Truncated protein isoforms generate diversity of protein localization and function in yeast

**DOI:** 10.1101/2023.07.13.548938

**Authors:** Andrea L. Higdon, Nathan H. Won, Gloria A. Brar

**Affiliations:** Department of Molecular and Cell Biology, University of California, Berkeley, Berkeley, CA 94720, USA; Center for Computational Biology, University of California, Berkeley, Berkeley, CA 94720, USA

## Abstract

Genome-wide measurements of ribosome occupancy on mRNA transcripts have enabled global empirical identification of translated regions. These approaches have revealed an unexpected diversity of protein products, but high-confidence identification of new coding regions that entirely overlap annotated coding regions – including those that encode truncated protein isoforms – has remained challenging. Here, we develop a sensitive and robust algorithm focused on identifying N-terminally truncated proteins genome-wide, identifying 388 truncated protein isoforms, a more than 30-fold increase in the number known in budding yeast. We perform extensive experimental validation of these truncated proteins and define two general classes. The first set lack large portions of the annotated protein sequence and tend to be produced from a truncated transcript. We show two such cases, Yap5^truncation^ and Pus1^truncation^, to have condition-specific regulation and functions that appear distinct from their respective annotated isoforms. The second set of N-terminally truncated proteins lack only a small region of the annotated protein and are less likely to be regulated by an alternative transcript isoform. Many localize to different subcellular compartments than their annotated counterpart, representing a common strategy for achieving dual localization of otherwise functionally identical proteins.

## INTRODUCTION

Defining the set of proteins encoded by an organism allows understanding of cellular function. This fundamental idea was the motivation for systematic coding sequence prediction immediately following the generation of whole genome sequences. Initial annotation strategies had known limitations, including difficulty in predicting splice isoforms and a reliance on strict rules to pare down the number of predicted open reading frames (ORFs), such as a minimum length requirement of 100 codons (reviewed in Dinger et al., 2008). Alternative splicing is generally considered to be the major force driving diversity of protein products from a single locus, but this process is relatively rare in budding yeast (although less rare than previously appreciated) (Ares et al., 1999; Douglass et al., 2019). Therefore, a simple model in which one gene encodes one transcript which is then decoded into one protein product has been adopted as a general rule in this organism, with a few isolated exceptions. Genomic techniques, such as transcript isoform sequencing and ribosome profiling, have greatly expanded our understanding of the diversity of transcript and protein products encoded by even very compact genomes like that of budding yeast (Brar et al., 2012; Chia et al., 2021; Eisenberg et al., 2020; Ingolia et al., 2011; Pelechano et al., 2013). Ribosome profiling, in particular, has facilitated the identification of a diverse array of translated open reading frames (ORFs), including upstream open reading frames (uORFs), intergenic short ORFs (sORFs), and N-terminally extended protein isoforms.

Meiosis in budding yeast is an excellent system for identifying fundamental principles of genome decoding and regulated gene expression. The process of differentiation from a diploid progenitor into haploid gametes requires an intricate and precisely timed series of cellular remodeling events (reviewed in Marston & Amon, 2004; van Werven & Amon, 2011). Underlying these dramatic changes is a gene expression program that requires dynamic regulation of most of the yeast proteome (Brar et al., 2012). Ribosome profiling and transcription start site sequencing show that production of non-canonical protein products and alternative transcripts is particularly prevalent during meiosis relative to mitotic growth (Brar et al., 2012; Chia et al., 2021; Sing et al., 2022). The specific functional relevance of all but a few non-canonical gene products, however, remains unclear. N-terminally truncated proteins, which initiate from downstream in-frame start codons within annotated genes, are a particularly interesting class. Since they are variants of existing proteins, they would seem likely to have molecular function, but their identification has remained difficult, both by sequencing-based and proteomics methods, due to specific technical challenges arising from their complete overlap with annotated coding regions.

Here, we develop a novel algorithm and identify hundreds of N-terminally truncated isoforms using data from a modified version of ribosome profiling for identifying translation initiation sites. In addition, we describe two distinct types of regulation responsible for their production, give experimental evidence for their existence, and provide insights into their functions. A handful of N-terminally truncated proteins have been validated in several organisms, including humans, and ribosome-profiling-based analyses suggest they might be common in mammalian cells (Ingolia et al., 2011). Although several individual examples of N-terminally truncated proteins were identified in previous single-gene studies in yeast (Table 1), our genome-wide approach, and subsequent experimental validation, sheds new light on their prevalence in this heavily studied model organism. Most of the 388 truncations we identified are dynamically regulated during the meiotic program or under other non-standard laboratory growth conditions. The compactness of the yeast genome and minimal use of splicing have led to the presumption that yeast do not widely use alternative protein isoforms. In fact, these very features facilitated our robust global analyses and investigation of their production.

**Table 1.**
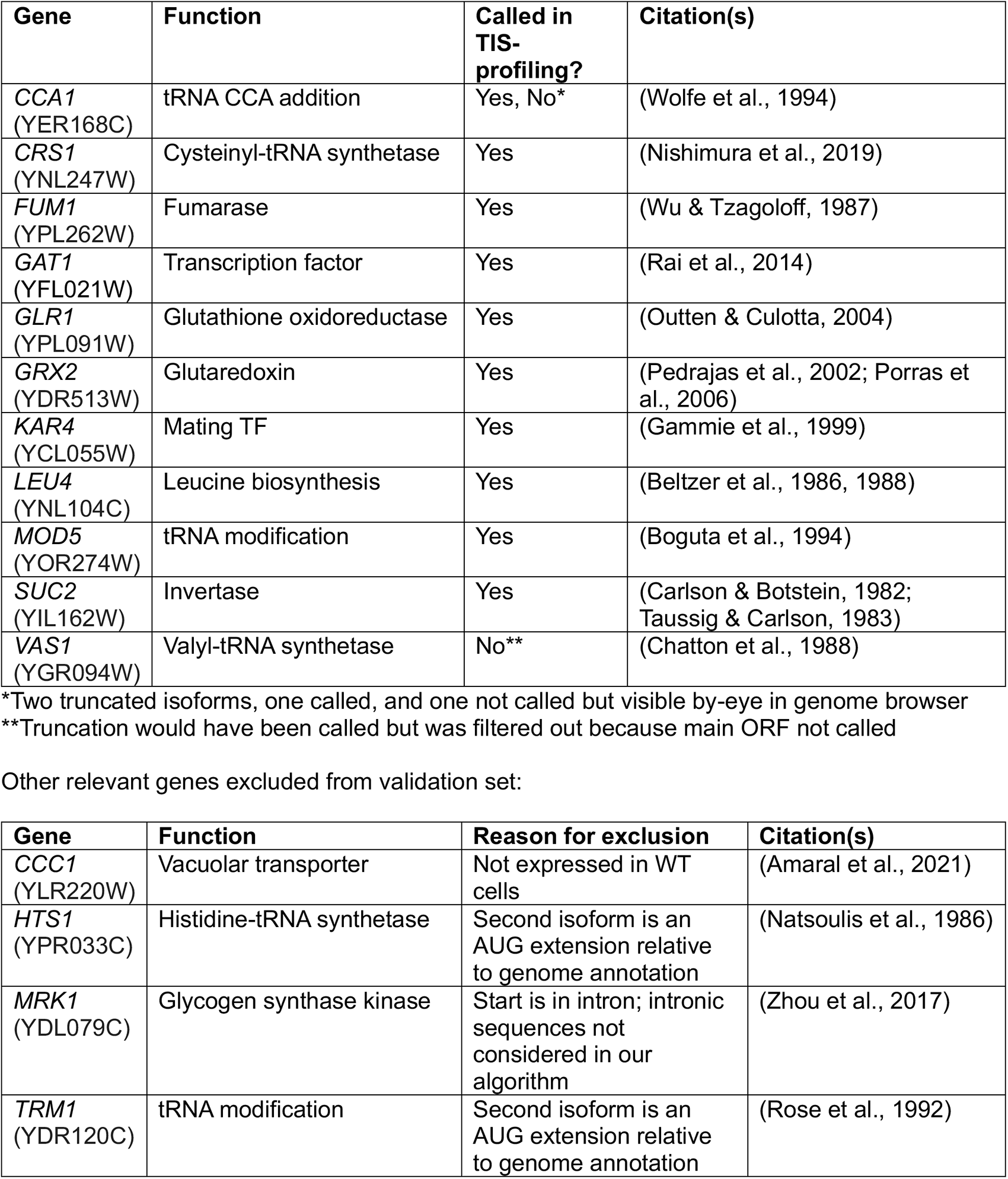
Previously characterized genes with N-terminally truncated protein isoforms.

We classified truncations into two broad classes that generally differentiate regulatory and functional characteristics of the isoforms. The first class of truncations lack a large portion of the annotated protein’s N-terminus (“distal” truncations) and tend to be encoded by a truncated transcript isoform. We identify and characterize such two examples, produced from the *PUS1* and *YAP5* loci, whose conserved and well-characterized annotated isoforms encode a pseudouridine synthetase and an AP-1 transcription factor, respectively. The truncated isoforms are expressed in a condition-specific manner and lack key domains, resulting in functions that seem distinct from the full-length proteins. The second class of truncations begin closer (“proximal”) to the annotated start codon. Proximal truncations are more often encoded by the same transcript as the annotated isoform, likely requiring bypass of the annotated start codon for their translation and allowing simultaneous production of the annotated and truncated isoform from a single transcript. Based on our extensive computational and experimental investigation, we posit that a common role for these truncations is in diversifying the subcellular localization of the encoded protein. We demonstrate that our predictions in this respect are remarkably robust, revealing a case in which two truncations at one locus allow three distinct and simultaneous subcellular localizations. Thus, our study elucidates the potential of truncated protein isoforms to provide proteome diversity and cellular function beyond what was previously recognized.

## RESULTS

### Truncated protein isoforms are prevalent in budding yeast

N-terminally truncated protein isoforms initiate at in-frame start codons within canonical genes and therefore share common C-terminal sequence with their annotated isoform. These truncated isoforms are more challenging to identify than many other types of translated regions. Standard ribosome profiling data, which have enabled global maps of translated regions, report the positions of elongating ribosomes and therefore yield reads mapping across the entire gene, including the entirety of the potential truncated isoform, masking signal for truncated isoforms (Figure 1A). We previously published a translation-initiation site (TIS-) profiling dataset with timepoints spanning stages of meiotic progression in budding yeast cells with the goal of identifying all types of non-canonical start sites, with a particular focus on N-terminally extended protein isoforms (Eisenberg et al., 2020). In contrast to standard ribosome profiling, which employs the translation elongation inhibitor cycloheximide (CHX), TIS-profiling uses the drug lactimidomycin (LTM) to capture ribosomes immediately after initiation, while allowing elongating ribosomes to complete translation (or “run off”; Figure 1A) (Ingolia et al., 2011; Lee et al., 2012). The resulting ribosome footprints are therefore highly enriched at sites of translation initiation, with minimal background reads across the ORF. TIS-profiling therefore yields clear start site peaks that are much easier to detect than with standard ribosome profiling. For example, at the *MOD5* locus, we detect both a previously identified truncated isoform (Mod5^truncation1^) and an additional previously unidentified truncated isoform (Mod5^truncation2^) using TIS-profiling (Figure 1B). Notably, neither truncated isoform is evident from standard ribosome profiling data, demonstrating the power of this method for detecting internal translation start sites.

**Figure 1.**
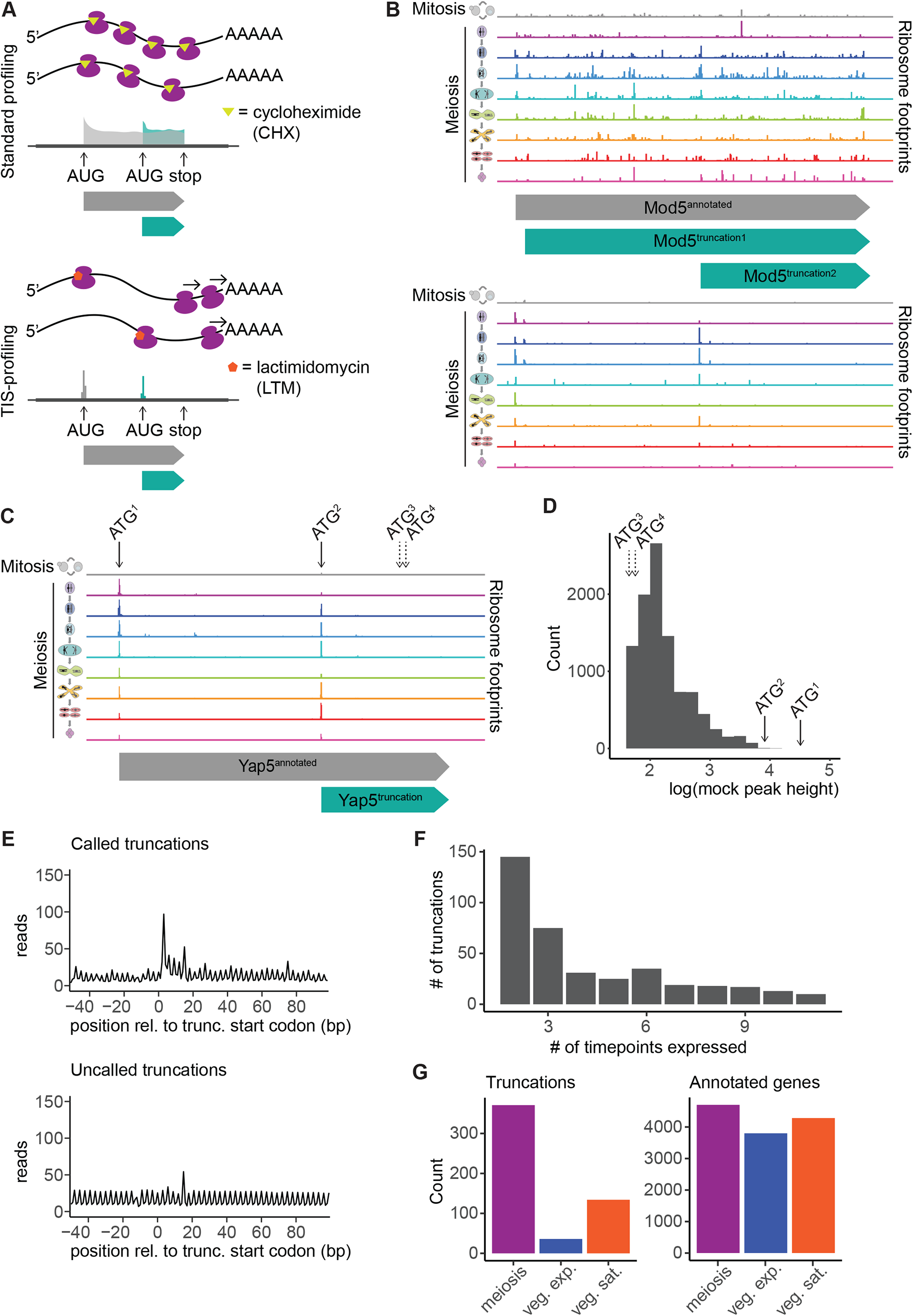
Genome-wide identification of truncated protein isoforms using TIS-profiling data. A. Schematic comparing standard ribosome profiling (top) and TIS-profiling (bottom). In each case, a cartoon of ribosomes translating two mRNAs is shown on top and sample read density is shown below for a locus containing an annotated and truncated protein isoform. B. Standard ribosome profiling (top) and TIS-profiling (bottom) data for the *MOD5* locus, with the annotated open reading frame in gray and the two truncations identified by our algorithm in turquoise (middle). Cartoons to the left indicate whether the track represents mitotic cells (gray) or meiotic timepoints (rainbow). Note that the two truncations are not apparent from the standard ribosome profiling data but robust peaks at multiple timepoints are clear in the TIS-profiling data. C. TIS-profiling data at the *YAP5* locus. Arrows indicate all in-frame ATGs within the ORF. Cartoons to the left indicate whether the track represents mitotic cells (gray) or meiotic timepoints (rainbow). D. Empirical distribution representing background read density within the *YAP5* annotated ORF at a single TIS-profiling time point (0h). Arrows indicate approximate read density for called start sites (solid arrows, ATG^1^ and ATG^2^) and uncalled in-frame start codons (dashed arrows, ATG^3^ and ATG^4^). E. Metagene plot of standard ribosome profiling data for all called truncated isoforms (top) and uncalled controls (bottom) for the region between -50 and +100nt relative to the truncation start codon. Reads are summed across all timepoints. For called truncations, downstream (+20 to +70bp) read density is significantly higher than upstream (-50 to -1bp) read density (p<0.01, Mann-Whitney U Test). For uncalled truncations there was not a significant difference in upstream and downstream read densities (n.s., Mann-Whitney U Test). F. Histogram of number of timepoints at which truncated isoforms are expressed in TIS-profiling data. G. Number of truncated isoforms (left) and number of annotated isoforms (right) called in samples from meiotic timepoints, vegetative exponential growth, and vegetative saturated growth. The variation in the number of called isoforms between conditions is significantly more pronounced for truncated isoforms than for annotated isoforms (p<2.2e^-16^, Fisher’s Exact Test).

We previously used the program ORF-RATER to call all types of open reading frames, including those that are non-canonical, using standard ribosome profiling and TIS-profiling data collected in vegetative (mitotic exponential and saturated) conditions and 8 timepoints spanning the major developmental stages of meiosis (Eisenberg et al., 2020; Fields et al., 2015). Although this algorithm performed very well for many types of ORFs, including those that were 5’ extended, it was unable to identify many truncated isoforms that were extremely clear in the start-site profiling data, such as at the *YAP5* locus (ATG^2^, Figure 1C) (Fields et al., 2015). We hypothesized that this could be due to overweighting of the standard ribosome profiling and underweighting of the TIS-profiling data by the algorithm. It has also been observed that many ribosome profiling algorithms that are trained on annotated ORFs are unable to perform as well with shorter ORFs (Spealman et al., 2021). To avoid systematically biasing against sensitive detection of translation initiation sites within annotated ORFs, we developed a novel algorithm specifically designed for identifying truncated isoforms that relies solely on the TIS-profiling data.

Briefly, the goal of this algorithm is to robustly interpret TIS-profiling data by separating true start-site signal from the background noise present across translated genes, likely due to low levels of elongation inhibition by LTM. To model the background signal for each gene at each timepoint, we randomly sampled three single nucleotide positions from within the annotated open reading frame and summed their mapped reads to simulate the three nucleotides of a start codon, over which a prominent peak should be present for real translation initiation sites. Resampling 10,000 times provided an empirical null distribution representing the distribution of peak heights that would be expected from the background noise within each gene at each time point (as shown for the *YAP5* locus at 0h in SPO, Figure 1D). We next determined the observed peak height for all in-frame (potentially truncation-generating) start codons within each annotated gene and used the corresponding empirical distribution to assign a p-value to each putative truncation start site. In the case of *YAP5*, there were three in-frame start codons within the annotated ORF, two of which fall within the range of peak heights expect from background and are not called (ATG^3^ and ATG^4^), and one which was called as significantly above background (ATG^2^) (Figure 1C-D). To increase the stringency of our calls, we required that a translation initiation site be called at 2 or more timepoints. We also required that a truncated isoform begin 5 amino acids (aa) or more from the annotated start codon and be no less than 10aa in total length. Truncated isoforms for which we could not detect the annotated isoform at any time point were also excluded, as these often represent cases of misannotation, where the “truncated” isoform is in fact the main isoform. Additional filtering criteria were applied, as described in the methods.

Using this approach, we identified 388 truncated protein isoforms. While the existence of truncated protein isoforms has been known for decades due to observations from single-gene studies, this represents a substantial increase in the number of known cases. To our knowledge, in *S. cerevisiae*, 12 truncated isoforms (two of which are present at the same locus, *CCA1*) have been characterized that we would expect to be present in our dataset as well (Table 1). Of these previously known truncations, 10 (83.33%) were called by our algorithm. Of the two that were not called, one was called by the algorithm but filtered out because its annotated isoform was not called (at the *VAS1* locus). The other (the shortest isoform of *CCA1)* was visible by eye in the TIS-profiling data but was not called by our algorithm. We concluded that our approach was sensitive to identifying translation of truncated isoforms and that they are much more prevalent than previously known, with our set of newly identified truncations representing a more than 30-fold increase over the set of characterized truncations.

Since our algorithm used only TIS-profiling data to identify truncated isoforms, we were able to evaluate the quality of our calls using matched standard ribosome profiling data. In these data, we would expect each truncated isoform to have a peak in reads at the start site, followed by slightly elevated read density downstream relative to upstream, since downstream read density should include the contributions of elongating ribosomes associated with both the annotated and truncated isoforms (Figure 1A). We performed metagene analysis for the regions surrounding the predicted start codon for all 388 truncations and indeed saw the expected read density patterns across the gene set (Figure 1E, upper). Read densities downstream of the start codon (+20 to +70bp; excluding the non-quantitative region immediately following the start codon) were higher those upstream (-50bp to -1bp), consistent with elevated ribosome footprint density corresponding to the translation of truncated isoforms (p<0.01, Mann-Whitney U Test). Importantly, this trend was not seen for in-frame start codons that were not called by our algorithm (n.s., Mann-Whitney U Test; Figure 1E, lower). Metagene profiles of the TIS-profiling data also showed the expect trends, with a sharp peak present at the start codon for called truncations, and virtually no signal at the start codon for uncalled truncations (Figure S1A-B). Together this indicates that our truncation calling approach detects true translation events.

### Truncated isoforms are dynamically expressed and enriched in meiosis

Among our set of called truncated isoforms, we observed that many appear to be dynamically expressed during meiosis. At the *MOD5* locus, for example, the smaller truncation (Mod5^trunc.2^) is not present in exponentially growing mitotic cells but is upregulated specifically during early meiosis (Figure 1B). Over 50 percent of truncated isoforms in our dataset were called at only two or three timepoints, indicating that dynamic expression is very common (Figure 1F). Truncated isoforms were also much more common in meiosis and slightly more common in vegetative saturated growth than in vegetative exponential growth, the most common laboratory growth condition (Figure 1G). Since we analyzed multiple samples collected during meiotic progression but only one each during vegetative exponential growth and vegetative saturated growth, this enrichment could have been due to increased power to detect ORFs during meiosis. However, this pattern was significantly less strong among annotated protein isoforms called by our algorithm, indicating that dynamic meiotic expression of truncated protein isoforms is a true biological phenomenon (p<2.2e^-16^, Fisher’s Exact Test).

### Newly predicted truncated protein isoforms can be detected *in vivo*

To evaluate the quality of our TIS-profiling data and truncation-calling algorithm, we experimentally validated the production of several newly identified truncated proteins. Due to the limitations in resolving similarly sized proteins, we focused on candidates that differed enough in size to be distinguishable from their annotated isoform by western blotting. We integrated a C-terminal epitope-tag at the endogenous locus of each protein, such that both the annotated and truncated isoforms should be tagged. We then collected samples at timepoints throughout meiosis and performed western blotting. Ten predicted truncated isoforms that we examined were clearly detectable by this method, indicating that the truncated proteins are expressed and stable. Some of these truncated protein isoforms display dynamic expression patterns distinct from their annotated isoform, including Yap5^trunc.^, Mod5^trunc.^, Pex32^trunc.^, Pus1^trunc.^, Sas4^trunc.^, Ssp1^trunc.2^, Tpo1^trunc.^, and Prp4^trunc.^ (Figure 2A-F, Figure S2A-B). Others, like Ssp1^trunc.1^ and Yck1^trunc.1^, display expression patterns that mirror the annotated isoform (Figure 2F-G).

**Figure 2.**
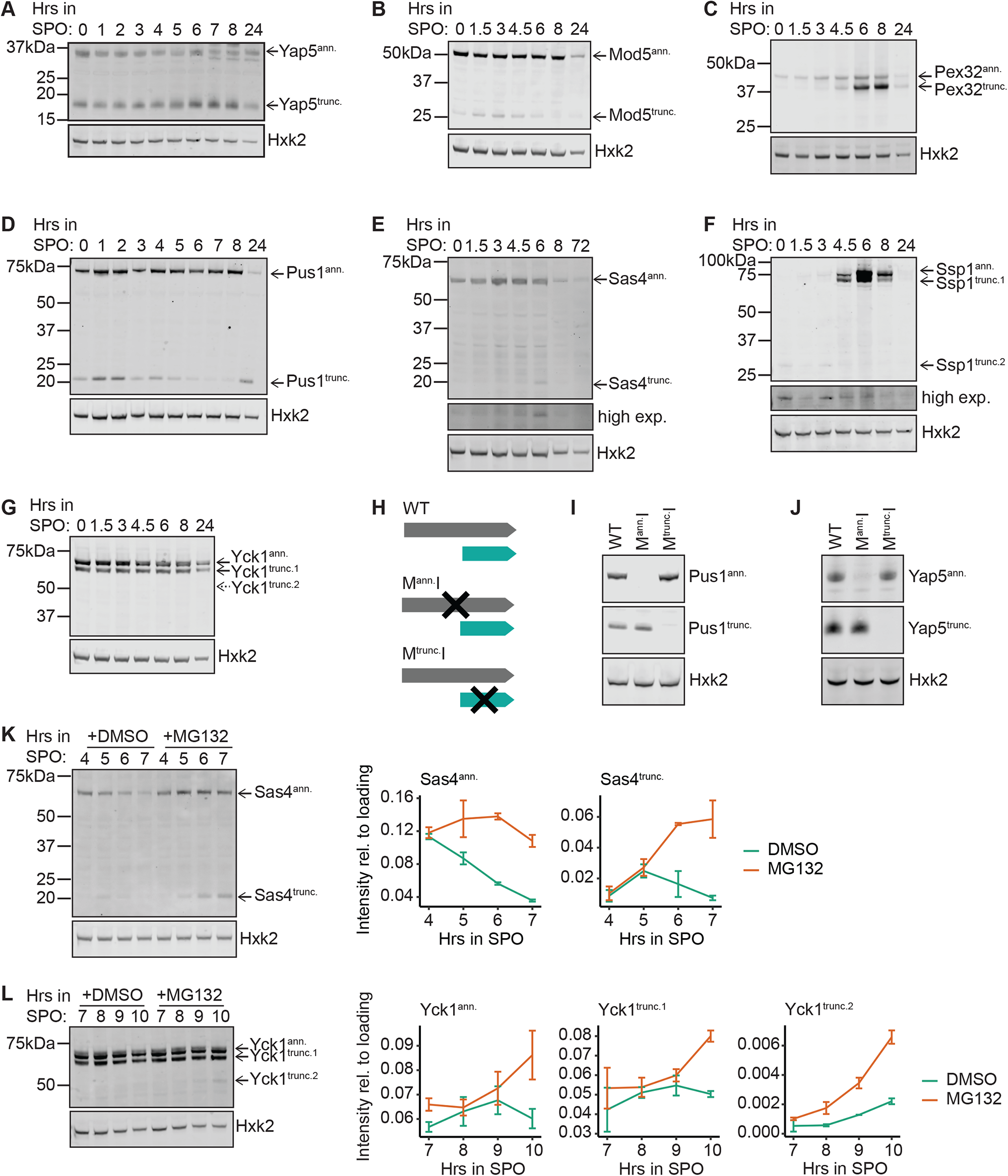
Many newly identified truncated protein isoforms can be confirmed by western blotting, some are stabilized by proteasome inhibition. A. Western blot of samples collected at various timepoints after transfer of cells to sporulation media (SPO). Hexokinase (Hxk2) is shown as a loading control. For lowly expressed truncated isoforms, a high exposure panel is shown (high exp.). A different strain is used in each case, expressing the indicated C-terminal epitope-tagged protein and enabling detection of annotated (ann.) and truncated (trunc.) isoforms for (A) Yap5-FLAG, B. (B) Mod5-3V5, C. (C) Pex32-3V5, D. (D) Pus1-3V5, E. (E) Sas4-3V5, F. (F) Ssp1-3V5, for with two truncated isoforms were predicted and detected, G. and (G) Yck1-3V5, for which two truncated isoforms were predicted but only one was successfully detected. Solid arrows indicate isoforms that were predicted and detected. Dashed arrows indicate predicted but undetected isoforms. H. Schematic of mutagenesis approach used to validate production of truncated isoforms from predicted start codons in (I) and (J). Top image indicates the wild-type context (WT), in which both isoforms should be seen. Either the annotated (middle, M^ann.^I) or predicted truncation (bottom, M^trunc.^I) start codon were mutated from encoding methionine (ATG) to isoleucine (ATT) to prevent their translation. I. Western blot of start codon mutant strains described in (H). Hexokinase (Hxk2) is shown as a loading control. Images of bands for the annotated and truncation isoforms are from the same blot. Blotting is for Pus1-3V5. Cells were collected at 24h post-dilution in YEPD. J. Western blot of start codon mutant strains described in (H) for Yap5. Hexokinase (Hxk2) is shown as a loading control. Images of bands for the annotated and truncation isoforms are from the same blot. Blotting is for Yap5-FLAG. Cells were collected at 4.5h in SPO. K. Representative western blot (left) and quantification (right) for cells treated with proteasome inhibitor MG132 or vehicle control DMSO, showing stabilization of both the truncated and annotated isoforms of Sas4-3V5 for cells in meiosis. Note that quantification is based on 2 replicates and error bars represent standard error. Hexokinase (Hxk2) is shown as a loading control. L. Representative western blot (left) and quantification (right) for cells treated with proteasome inhibitor MG132 or vehicle control DMSO, showing stabilization of the truncated and annotated isoforms of Yck1-3V5 for cells in meiosis. Note that quantification is based on 2 replicates and error bars represent standard error. Hexokinase (Hxk2) is shown as a loading control.

Although the detected truncated isoforms migrate according to their expected size, it remained possible that the bands could be degradation products of the annotated isoform rather than the product of translation at the predicted alternative in-frame start codon. To test this possibility, we generated strains for two examples, Pus1 and Yap5, for which the ATG start codons for the annotated or predicted truncated isoform were mutated to ATT (isoleucine) to abrogate expression (M^ann.^L and M^trunc.^L, respectively; Figure 2H). Western blot analysis of these strains revealed translation of only the annotated isoform in cells carrying the M^trunc.^L mutation, and only the truncated isoform in cells carrying the M^ann.^L mutation, indicating that the truncated isoforms for each gene are indeed the product of translation initiating at the newly predicted start codon internal to the annotated ORF (Figure 2I-J).

Our ability to detect most of the predicted truncated isoforms that we tested suggest our approach has a low rate of false positives. However, 5 out of 15 predicted truncations that we tested were not visible by western blotting (Yck1^trunc.2^, Glk1^trunc.^, Ari1^trunc.^, Rtt105^trunc.^, and Siw14^trunc.^; Figure 2G, Figure S2C-F). There are several possible explanations for this: (1) the truncated protein may not be compatible with the specific epitope tag, (2) they may be challenging to detect for technical reasons, for example due to low expression or small size, (3) they could be false positives, or (4) they may be produced but then degraded under normal conditions. To test whether protein degradation was preventing our detection of some truncated protein isoforms, we treated cells carrying C-terminal epitope tags of predicted truncations with the proteasome inhibitor MG132. We timed MG132 treatment and sample collection for western blotting according to the expected timing of expression for each truncation based on the TIS-profiling data. We included examples in which the truncation was not detected (Yck1^trunc.2^, Ari1^trunc.^, Rtt105^trunc.^, and Siw14^trunc.^) as well as ones in which the truncation was visible but present at low abundance (Sas4^trunc.^, Tpo1^trunc.^, Mod5^trunc.^, and Prp4^trunc.^) to look for evidence of stabilization. We indeed saw increased abundance, indicating increased stabilization, for previously detectable truncations, including Sas4^trunc.^ and Tpo1^trunc.^ (Figure 2K, Figure S2G). Interestingly, the previously undetectable second truncation of Yck1^trunc.2^ became visible upon proteasome inhibition, suggesting that this truncation is normally translated but degraded by the proteasome to levels below detection (Figure 2L). Abundance of other truncated proteins, including Mod5^trunc.^ and Prp4^trunc.^ (Figure S2H-I), was minimally affected by proteasome inhibition, suggesting that proteasome-mediated degradation was not the reason for their weak detection by western blotting. Three of the previously undetectable truncations (Ari1^trunc.^, Rtt105^trunc.^, and Siw14^trunc.^) remained undetectable upon proteasome inhibition (Figure S2J-L).

### “Distal” truncations are typically produced from a truncated transcript while “proximal” truncations are likely regulated by translational control

Many truncated isoforms are dynamically regulated, some in concert with their annotated isoform and others independently. There are two straightforward models to explain such regulation: either (1) translational control, in which the ribosome bypasses one or more in-frame start codons in favor of a downstream one or (2) transcriptional control, in which a truncated transcript encodes the truncated protein and translation initiation occurs at the first in-frame start codon (Figure 3A). To determine the extent of these two possible types of regulation for the hundreds of new truncated proteins identified by our algorithm, we leveraged a published transcript leader sequencing (TL-seq) dataset collected across meiosis in the same strain background as our TIS-profiling data (SK1) (Chia et al., 2021). Using these data, we were able to determine whether there is evidence of a 5’ transcript end upstream of our called truncation TISs but downstream of their respective annotated TISs. For *YCK1*, for example, two truncated isoforms (Yck1^truncation1^ ^and^ ^2^) are produced but there is no evidence of a separate transcript isoform in the TL-seq data, consistent with translational control (Figure 3B). For *YAP5*, in contrast, a clear TSS is present just upstream of the TIS for the truncated isoform, supporting the transcriptional control model (Figure 3C).

**Figure 3.**
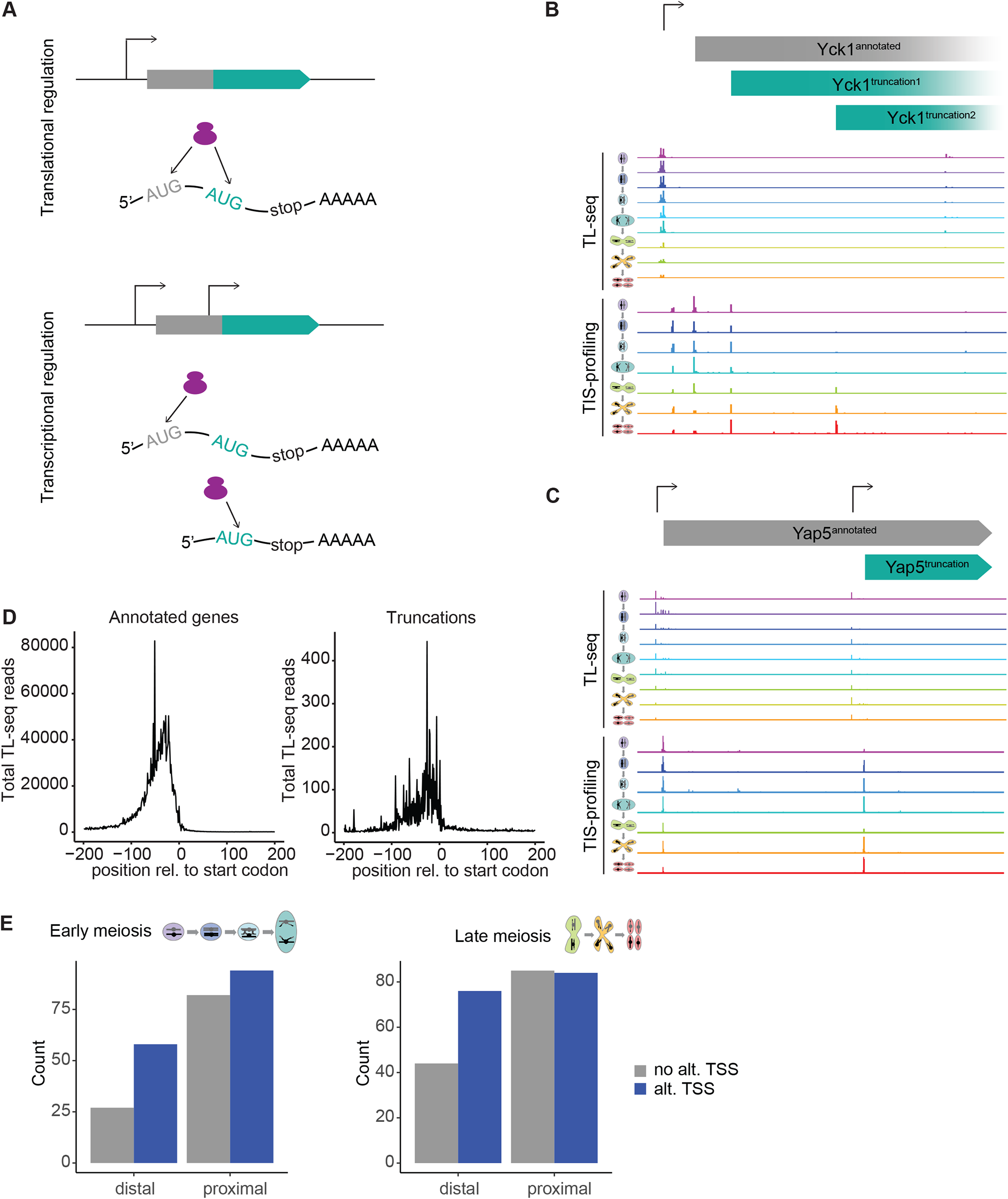
Truncated protein isoforms are often, but not always, produced from truncated transcript isoforms. A. Schematic of potential regulatory mechanisms for truncated protein isoform production: (1) Translational regulation (top) in which a single transcript isoform is produced, and protein isoform production is determined by initiation site selection. (2) Transcriptional regulation (bottom) in which an annotated and truncated transcript isoform are translated into annotated and truncated protein isoforms, respectively. In each case, a schematic of the locus is shown with bent arrows representing transcription start sites above and resulting transcript(s) shown below. B. TL-Seq (above) and TIS-profiling (below) data for the *YCK1* locus showing production of the annotated isoform and two truncated protein isoforms but evidence for only the canonical transcript isoform. Cartoons at left represent meiotic timepoints (rainbow). Bent arrow represents the predicted transcription start site. The start peak upstream of the annotated start codon corresponds to an upstream open reading frame (uORF). C. TL-Seq (above) and TIS-profiling (below) data for the *YAP5* locus showing annotated and truncated transcript and protein isoform production. Cartoons at left represent meiotic timepoints (rainbow). Bent arrows represent predicted transcription start sites. D. Metagene plots of total TL-seq reads for the window -200 to +200bp surrounding translation initiation sites for annotated (left) and truncated (right) protein isoforms. Truncations with an annotated start beginning within the -200bp window were excluded to avoid including reads derived from annotated transcripts. E. Bar plot of number of distal (>40aa from annotated start) and proximal (≤40aa from annotated start) truncated isoforms with evidence of an alternative transcription start site (“alt. TSS”) or not (“no alt. TSS”) based on interpretation of 5’ ends in approximately stage-matched TL-seq data collected in either early (left) or late (right) meiosis. Distal isoforms are significantly more likely to show evidence of an alternative transcript isoform (p<0.05 for both early and late groupings, Fisher’s Exact Test).

To assess the prevalence of truncated transcripts corresponding to truncated proteins genome-wide, we constructed TL-seq metagene profiles for the regions upstream of translation initiation sites (Figure 3D). For annotated genes, this displays the expected profile of high read density peaking around 50bp upstream of the TIS, consistent with the average length of a yeast 5’ UTR (Nagalakshmi et al., 2008). The metagene for truncated isoforms displays a strikingly similar profile, indicating that it is common for truncated proteins to be produced from truncated transcripts (Figure 3D). Importantly, for this analysis we excluded truncated isoforms whose annotated isoform would start within the window included in the metagene profile (-200bp) such that the metagene profile should not include any reads derived from annotated transcript isoforms.

We hypothesized that regulation via truncated transcripts might be more common for truncations that initiate far (distal) from the annotated isoform since control at the translational level would in most cases require the ribosome to bypass several in-frame start codons, making transcriptional regulation a more parsimonious explanation. Truncated isoforms starting close (proximal) to the annotated start site, however, would be more likely to only require bypassing of the annotated start codon which could easily occur due to leaky scanning (Kozak, 2005). For purposes of this comparison, we defined “proximal” truncations as starting within 40aa of the annotated start codon and “distal” truncations as starting greater than 40aa from the annotated start codon. To determine which truncated isoforms had evidence of a corresponding truncated transcript, we used TL-seq data to calculate a “TSS score” which is the ratio of the sum of TL-seq reads 200bp upstream of the truncation TIS over the sum of reads 200bp downstream of the truncation TIS. A higher ratio indicates stronger evidence of an alternative transcription 5’ end (likely generated by a downstream transcription start site) that is close to the truncation TIS. To assess statistical significance, we compared each ratio to an empirical distribution of TSS scores created by randomly sampling 200bp windows from within the same gene. Due to the differences in staging between the two meiotic time courses, we were unable to do high-resolution time point matching between the time courses. However, we were able to achieve a level of temporal resolution that was still compatible with the differences in staging by splitting timepoints into either early or late meiosis based on correlation coefficients between time points in matched mRNA-seq data for each time course, as well as expression patterns of key meiotic genes (Figure S3A-B). From this analysis we observed that distal truncations were much more likely to have a TSS than proximal truncations, with approximately two-thirds of distal truncations showing evidence of transcriptional regulation via a truncated transcript (p<0.05 for both early and late meiotic groups, Fisher’s Exact Test, Figure 3E).

### Distal truncated protein isoforms for Yap5 and Pus1 exhibit condition-specific regulation

To better understand the functional relevance of distal, transcriptionally regulated truncations, we performed in-depth characterization of two examples, at the *YAP5* and *PUS1* loci. TIS-profiling data showed a very sharp start peak at an in-frame start codon within the *YAP5* locus, indicating translation of a truncated protein isoform (Yap5^truncation^) across all meiotic time points (Figure 1C). Western blot analysis confirmed that the truncation is expressed across meiosis but is low during vegetative exponential growth (Figure 2A, Figure 4A). Matched TL-seq data shows evidence of a transcript isoform with its 5’ end positioned approximately 30bp upstream of the Yap5^truncation^ start codon, suggesting that the truncated protein isoform is produced from an alternative transcript (Figure 3C). Given the dynamic regulation of Yap5^truncation^ we wondered whether it had a role related to that of the full-length protein. Annotated Yap5 is an iron-response transcription factor that upregulates its target genes upon exposure to elevated iron in order to mitigate iron toxicity (Li et al., 2008). It constitutively occupies its target promoters and activates target gene transcription upon binding to Fe-S clusters in high iron conditions. The DNA binding domain of Yap5 is in the N-terminal half of the protein and is not present in Yap5^truncation^; the Fe-S cluster binding domain lies in the C-terminal half and remains intact in the truncated isoform (Li et al., 2008). Previous work with artificial truncations of Yap5 showed that a version containing a comparable C-terminal region alone is capable of binding Fe-S clusters, suggesting that the natural truncation is also capable of this function (Rietzschel et al., 2015). We therefore hypothesized that Yap5^truncation^ may play a role in Fe-S cluster homeostasis.

**Figure 4.**
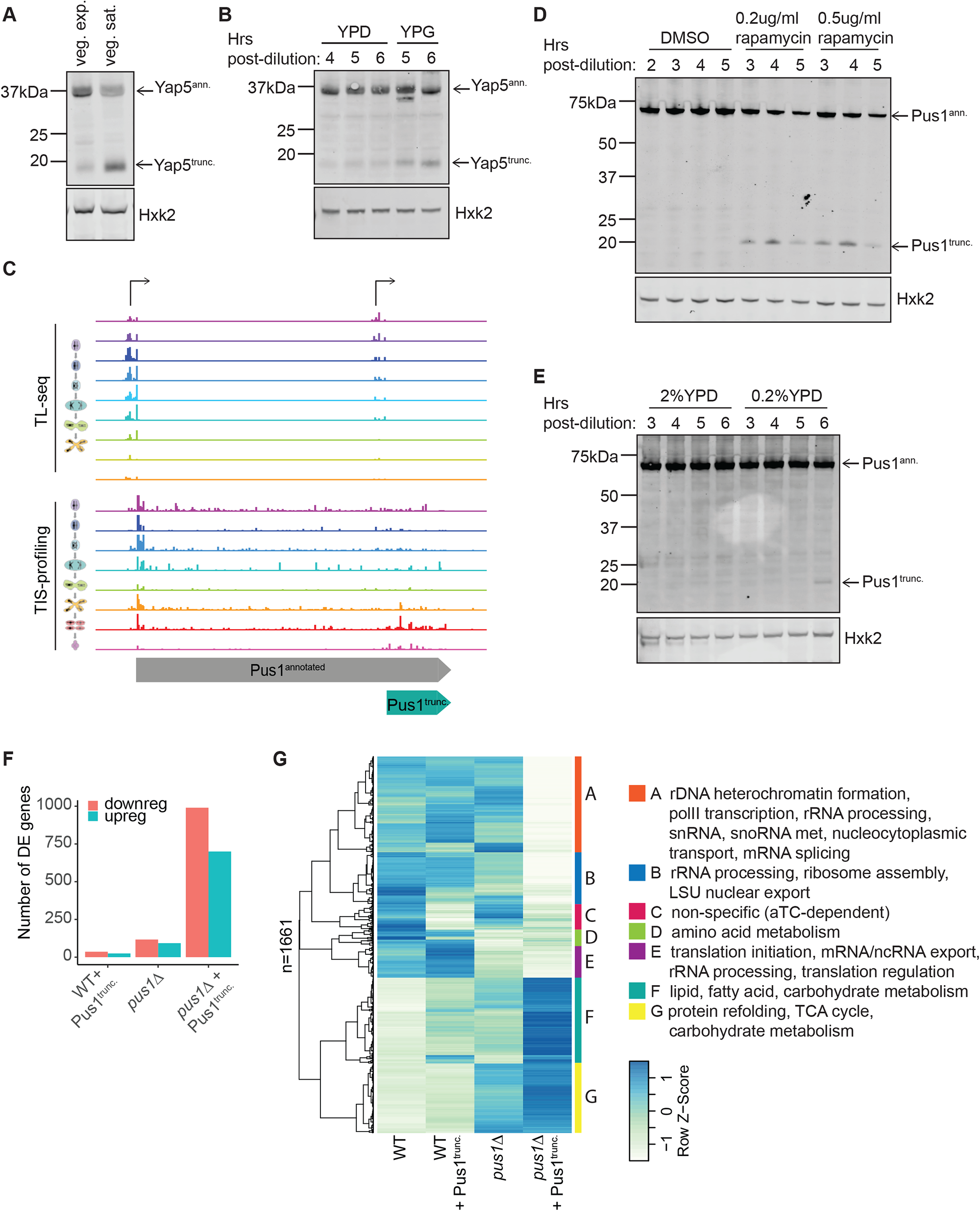
Condition-specific regulation of distal truncated protein isoforms. A. Western blot for Yap5-FLAG. Samples were collected during vegetative exponential and vegetative saturated growth in YEPD. Hexokinase (Hxk2) is shown as a loading control. B. Western blot for Yap5-FLAG from cultures grown in rich media (YEPD) for 4 hours then transferred to either YEPD (fermentable) or YEPG (non-fermentable) media. Hexokinase (Hxk2) is shown as a loading control. C. TL-seq (above) and TIS-profiling (below) data at the Pus1 locus showing annotated and truncated transcript and protein isoforms. Cartoons at left of each data type indicate meiotic timepoints (rainbow). Bent arrows represent predicted transcription start sites. Note that very early TL-seq timepoints are not fully stage-matched to the first TIS-profiling timepoint, and late TIS-profiling timepoints do not have matched equivalents in the TL-seq time course, leading to patterns that appear to differ in timing when comparing patterns between the two sets of samples. D. Western blot for Pus1-3V5 from samples grown in rich media (YEPD) and treated with either DMSO or two different rapamycin concentrations for the times indicated. Hexokinase (Hxk2) is shown as a loading control. E. Western blot for Pus1-3V5 from samples grown in either standard (2%YEPD) or reduced-nutrient (0.2%YEPD) media for the timepoints indicated. Hexokinase (Hxk2) is shown as a loading control. F. Bar graph showing the number of genes differentially expressed in the following conditions: WT cells carrying a construct to allow anhydrotetracycline-inducible Pus1^trunc.^ expression and treated with anhydrotetracycline (aTC; “WT + Pus1^trunc.^”), *pus1Δ* cells carrying a construct to allow aTC-inducible Pus1^trunc.^ expression and treated with vehicle (“*pus1Δ”*) or aTC (“*pus1Δ* + Pus1^trunc.^*”*). In all cases differential expression is relative to WT cells carrying a construct to allow aTC-inducible Pus1^trunc.^ expression and treated with vehicle (“WT”). Differentially expressed genes were determined using DESeq2. G. Clustered heatmap of RNA-seq data for all genes that were differentially expressed between WT cells treated with vehicle (“WT”) and *pus1Δ* cells treated with aTC (“*pus1Δ* + Pus1^trunc.^*”*). Samples include WT cells carrying a construct to allow aTC-inducible Pus1^trunc.^ expression and treated with vehicle (“WT”) or aTC (“WT + Pus1^trunc.^”), cells deleted for *PUS1* and carrying a construct to allow aTC-inducible Pus1^trunc^ expression and treated with vehicle (“*pus1Δ”*) or aTC (“*pus1Δ* + Pus1^trunc.^*”*). Values are the log of the average of two replicates. Seven discrete clusters are indicated to the right with colored bars and letters. Gene ontology terms enriched in each cluster are indicated to the right.

Fe-S clusters are key cofactors in the electron transport chain and thus important for mitochondrial function. It is therefore notable that Yap5^truncation^ is induced during meiosis, a condition requiring respiration, but is lowly expressed in mitotic growth conditions, which favor fermentation. We also observed increased Yap5^truncation^ expression in mitotic cells grown to saturation in rich media, a condition in which cells have undergone the diauxic shift from fermentation to respiration upon exhaustion of their fermentable carbon sources (Figure 4A). We wondered whether the Yap5 truncation would also be induced in other respiratory conditions. To test this, we grew cells in dextrose-containing (fermentable) media, and after 4 hours of growth split the cells into either dextrose- or glycerol-containing (non-fermentable) media. We then assayed Yap5^truncation^ production by western blotting and observed upregulation of Yap5^truncation^ upon the switch to non-fermentable media (Figure 4B). These experiments suggest that dynamic production of Yap5^truncation^ occurs as part of a cellular response to conditions of increased respiratory activity.

Experimental validation, as well as comparison to TL-seq and standard ribosome profiling data, support the robustness of our approach for identifying hundreds of new truncations, a class of non-canonical coding region that has been difficult to annotate sensitively and reliably. We chose parameters specifically to minimize false positive detection, a major issue for this class of protein in our experience (Eisenberg et al., 2020). However, this also necessitated excluding some promising predicted truncations that did not pass some of our filters. As an example, TIS-profiling data indicated that the *PUS1* locus also encodes two isoforms, the annotated isoform and a truncated isoform (Pus1^truncation^), but the data were noisier than cases like Yap5^truncation^, and this resulted in our algorithm only calling this truncation at one timepoint, leading to its exclusion from our final list of truncated isoforms (*YAP5:* Figure 3C, *PUS1:* Figure 4C). In support of Pus1^truncation^ representing a true case, TL-seq data additionally showed evidence of an alternative transcript isoform beginning about 50-70bp upstream of the Pus1^truncation^ start codon, indicating that the truncated protein is likely to be translated from an alternative transcript (Figure 4C). Moreover, Pus1^truncation^ was readily detected by western blotting of cells expressing a C-terminally 3V5-tagged Pus1, and is dynamically expressed across meiosis, with levels peaking early in meiosis and again later in spores (Figure 2D). After confirming expression of Pus1^truncation^, we next sought to understand its function. The annotated Pus1 protein is a pseudouridine synthase that is known to play key roles in translation through modification of tRNAs and snRNAs (reviewed in Rintala-Dempsey & Kothe, 2017). More recently, Pus1 has also been found to modify a subset of mRNAs (Basak & Query, 2014; Carlile et al., 2014; Lovejoy et al., 2014; Massenet et al., 1999; Motorin et al., 1998; Schwartz et al., 2014). Pus1^truncation^ contains regions involved in RNA binding, present at the C-terminus of the full-length protein, but lacks the N-terminal catalytic domain responsible for pseudouridylation in annotated Pus1 (Czudnochowski et al., 2013; Rintala-Dempsey & Kothe, 2017).

Our western blot data indicate that Pus1^truncation^ is most abundant in early meiosis and in spores, which are both times that display reduced levels of translation, as measured by polysome profiling (Brar et al., 2012). To assess whether decreased translation is associated with Pus1^truncation^ expression, we analyzed multiple additional conditions in which translation is reduced. First, we investigated the regulation of the *PUS1* locus in a standard ribosome profiling dataset collected in in meiotic cells lacking ribosomal protein Rpl40a, a condition which even more severely reduces translation early in meiosis (Cheng et al., 2019). Relative to WT, *rpl40aΔ* cells show reduced translation of the annotated Pus1 isoform and much more prominent translation of Pus1^truncation^ (Figure S4A).

To monitor Pus1^truncation^ levels in an orthogonal low-translation context, we treated mitotically growing cells with rapamycin, a general inhibitor of growth and ribosome biogenesis. Upon rapamycin treatment, we observed marked induction of Pus1^truncation^, compared to undetectable levels in the untreated vehicle control (Figure 4D). Glucose starvation is another condition that reduces translation. By western blotting, we compared Pus1^truncation^ production in cells grown in normal rich media (2% glucose YEPD) to those grown in low-glucose media (0.2% glucose YEPD). After several hours in low-glucose media, we observed slight induction of the truncated isoform, compared to no induction in standard rich media (Figure 4E). Together, these data are consistent with a role for Pus1^truncation^ in conditions in which cellular translation is lowered, perhaps also explaining why this truncation has not been observed previously in the many studies focused on standard nutrient-rich growth conditions.

To further understand the role of Pus1^truncation^, we strongly expressed the truncated isoform in vegetative exponential cells, a condition where the truncation is not normally present, and performed mRNA-seq to assess transcript level changes. We used an anhydrotetracycline-inducible allele to conditionally express Pus1^truncation^ and collected uninduced and induced samples in both a WT and deletion (*pus1Δ*) background (Figure S4B). Expression of Pus1^truncation^ alone had relatively little impact on overall gene expression, as measured by the number of differentially expressed genes relative to WT (n=61). Deletion of *PUS1* resulted in a slightly increased, but still modest, number of differentially expressed genes (n=210; p<2.2e^-16^, Fisher’s Exact Test). The combination of *pus1Δ* with Pus1^truncation^ expression, by contrast, yielded dramatically higher rates of differential gene expression, suggesting a synthetic phenotype between the two perturbations (n=1690; p<2.2e^-16^, Fisher’s Exact Test; Figure 4F, Figure S4C).

We performed gene ontology (GO) analysis of the differentially expressed genes in *pus1Δ* cells expressing Pus1^truncation^ and found that the genes that were downregulated in the *pus1Δ* with Pus1^truncation^ expression condition were strongly enriched for GO terms related to translation and RNA modification substrates, including ncRNA production and processing, rRNA processing, and ribosome biogenesis (Figure S4D). To better understand the nature of the synthetic phenotype and visualize expression patterns across all our samples, we performed hierarchical clustering of all genes that were differentially expressed between WT and *pus1Δ* with Pus1^truncation^ expression (Figure 4G). Cluster C contains genes that are upregulated in both WT and *pus1Δ* cells upon Pus1^truncation^ induction. This gene set is generally upregulated upon aTC-induction of entirely unrelated genes and is therefore likely to be drug-dependent rather than Pus1^truncation^-specific (data not shown). Cluster E contains genes that are strongly downregulated in *pus1Δ* cells but are unaffected by Pus1^truncation^ expression. This cluster is enriched for genes involved in translational regulation, RNA export, and rRNA processing, consistent with known roles of Pus1. In light of the observed synthetic effects of expression of Pus1^truncation^ in a *pus1Δ* background, two additional clusters are of particular note: (1) Cluster B shows very little change upon Pus1^truncation^ expression alone, modest downregulation occurring with the *pus1Δ* alone, and much more pronounced downregulation arising in the *pus1Δ* with Pus1^truncation^ expression. This cluster is enriched for rRNA processing and ribosome assembly. (2) Cluster A, in contrast, contains genes that are largely unaffected with either of the individual perturbations but are strongly downregulated in the *pus1Δ* with Pus1^truncation^ expression. This cluster is strongly enriched for genes involved in rDNA heterochromatin formation, rRNA processing, splicing, and snoRNA metabolism. Together these results indicate that, while the loss of Pus1 alone has some impact on processes related to translation and ribosome biogenesis, the addition of Pus1^truncation^ expression expands the number of affected genes and increases the severity of the changes to translation-related transcript expression. Thus, Pus1^truncation^ appears to have subtle but global effects on translation that manifest primarily in the sensitized hypomodified context of *pus1Δ* cells, arguing that the role of the truncated isoform of Pus1 is not dependent solely on the function of the annotated full-length Pus1 isoform but may be related to the functions of other pseudouridine synthases that affect translation.

### Proximal truncations are a general mechanism for encoding multiple localizations of protein products at a single locus

Although in-frame start codons are uniformly distributed throughout coding genes, their usage as translation start sites in our dataset is very non-uniform, with a strong bias for more N-terminal (proximal to annotated starts) start sites (Bazykin & Kochetov, 2011) (Figure 5A). In fact, nearly 60% of all truncations start within 40aa of the corresponding annotated start codon. We hypothesized that – unlike in the case of distal truncations, which are likely to encode isolated domains – for proximal truncations, the core functional domains of the protein should typically remain intact in the truncated isoform, but the missing N-terminal sequences could encode localization signals, resulting in differential localization of the truncated isoform. This hypothesis is supported by the fact that among the small set of previously known truncated isoforms in *S. cerevisiae*, nearly all serve the function of differentially localizing a subset of the protein (Table 1, Figure 5B). We reasoned that our dataset would be an excellent opportunity to test whether this is indeed a broad phenomenon and to potentially identify additional differentially localized truncated isoforms.

**Figure 5.**
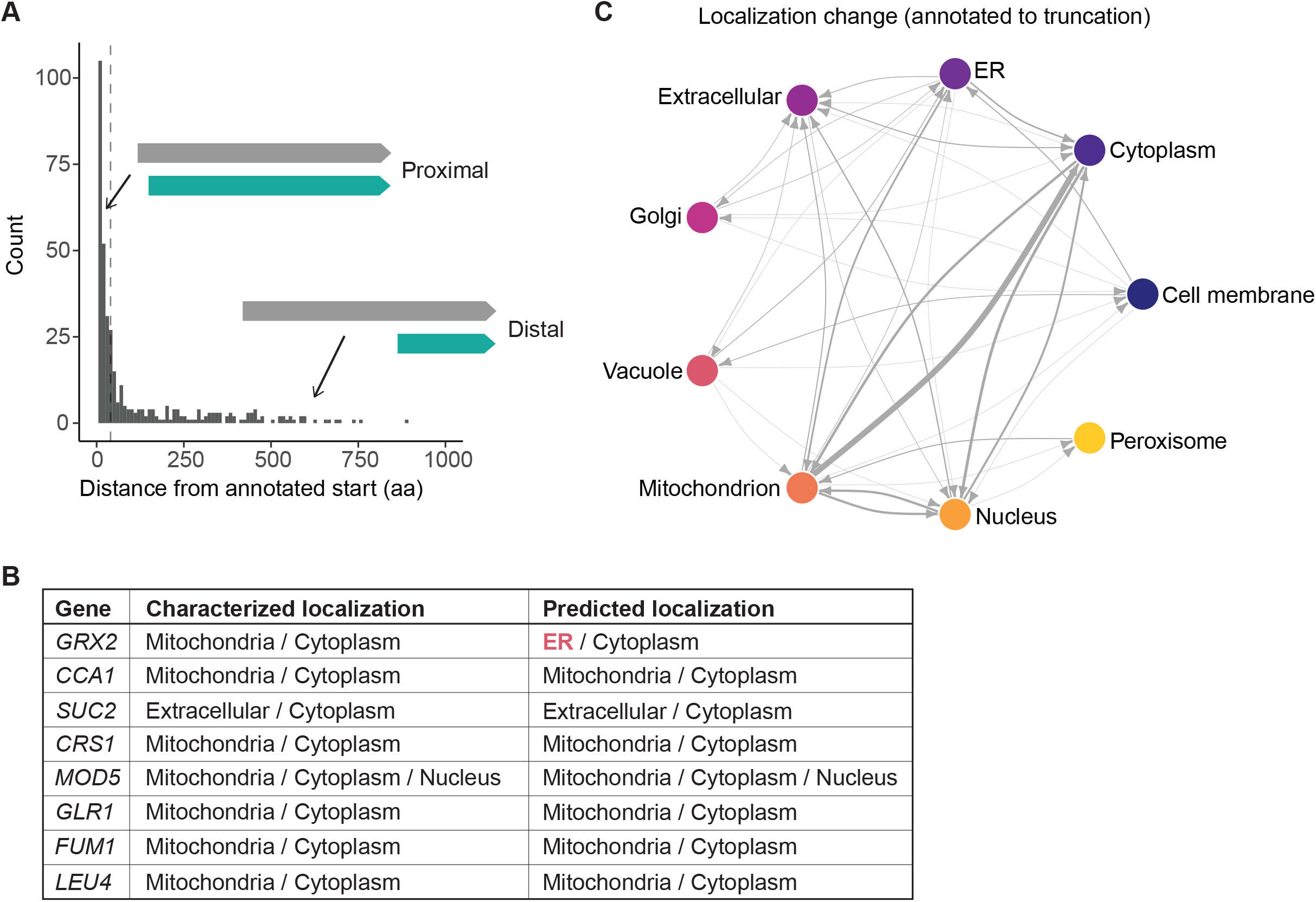
Computational prediction of differentially localized truncated isoforms. A. Distribution of distances between annotated and truncated isoform translation initiation sites among 388 truncations called by the algorithm, with cartoons above representing the “proximal” and “distal” categorization of truncations. The dotted line represents the 40aa cutoff between the proximal and distal categories. B. Table of previously characterized truncations called by our algorithm, comparing their experimentally characterized localization with their predicted localization using DeepLoc1.0. Discrepancies between predicted and characterized localization are indicated in red. Note that for one gene *CCA1*, our algorithm only called the larger of two known truncated isoforms, so the smaller isoform was not included in the localization predictions. C. Schematic of predicted subcellular localizations of annotated and truncated protein isoforms, with the blunt and pointed end of the arrows representing the annotated and truncated isoforms, respectively. Localization prediction was performed using DeepLoc1.0. Weight of line indicates number of truncations in each category.

We used a published algorithm, DeepLoc1.0, to perform localization prediction for our truncated isoforms and their annotated counterparts (Almagro Armenteros et al., 2017). DeepLoc1.0 is a deep learning algorithm trained on existing protein databases and uses sequence information alone to predict protein localization. It was important to use an algorithm that used sequence rather than homology, as the truncated and annotated isoforms would have very similar homology given that they share significant amounts of sequence. Since localization signals are frequently encoded at the N-terminus, we reasoned that “artificial” truncations could easily generate differentially localized proteins by removing an N-terminal sequence. To determine the expected background rate of differential localization we generated sets of simulated truncations by randomly sampling in-frame start codons and performing localization prediction on those open reading frames compared to the annotated protein. We found that our set of truncations differentially localized at a higher rate than expected by chance, with just over 30% of truncated isoforms localizing to a different compartment than their respective annotated isoforms (Figure S5A).

Among differentially localized truncations, it was most common for the truncations to lose the predicted mitochondrial or nuclear localization of their annotated counterpart, while cytoplasmic or extracellular localization were the most likely localizations to be gained (Figure 5C, Figure S5B). For initial validation of this prediction method, we compared the predicted and experimentally determined localizations for previously characterized truncations. For all but one of these genes, *GRX2*, the localization predictions match the characterized localization of the known isoforms, suggesting that the localization predictions were robust (Figure 5B).

We selected six predicted dual localized truncated proteins and sought to experimentally validate their localization using fluorescence microscopy (Figure S6A). Candidate proteins were primarily selected based on the quality of start-site profiling data at the relevant locus and their ability to be tagged and imaged. Five candidates had no known alternative isoforms in existing literature. The sixth, *GRX2*, had a characterized truncated isoform for which the localization predictions did not match the characterized localizations (Figure 5B). For each locus of interest, we inserted a fluorescently tagged transgene under the control of the gene’s endogenous promoter. To determine the localization of each isoform separately, we generated three strains for each gene: (1) all isoforms, with unmutated start codons, (2) annotated only, with the truncation start codon mutated to isoleucine, and (3) truncation only, with the annotated start codon mutated to isoleucine (Figure 6A, Figure S6B-D). Two genes, *IGO2* and *APE4*, were not possible to validate – the truncated isoform of Igo2 was not detectable by western blotting or microscopy, and although both isoforms of Ape4 could be detected by western blotting they were too low-abundance to detect by microscopy. For the remaining candidates, we performed fluorescence microscopy, collecting images at representative meiotic time points at which the respective truncations are expressed.

**Figure 6.**
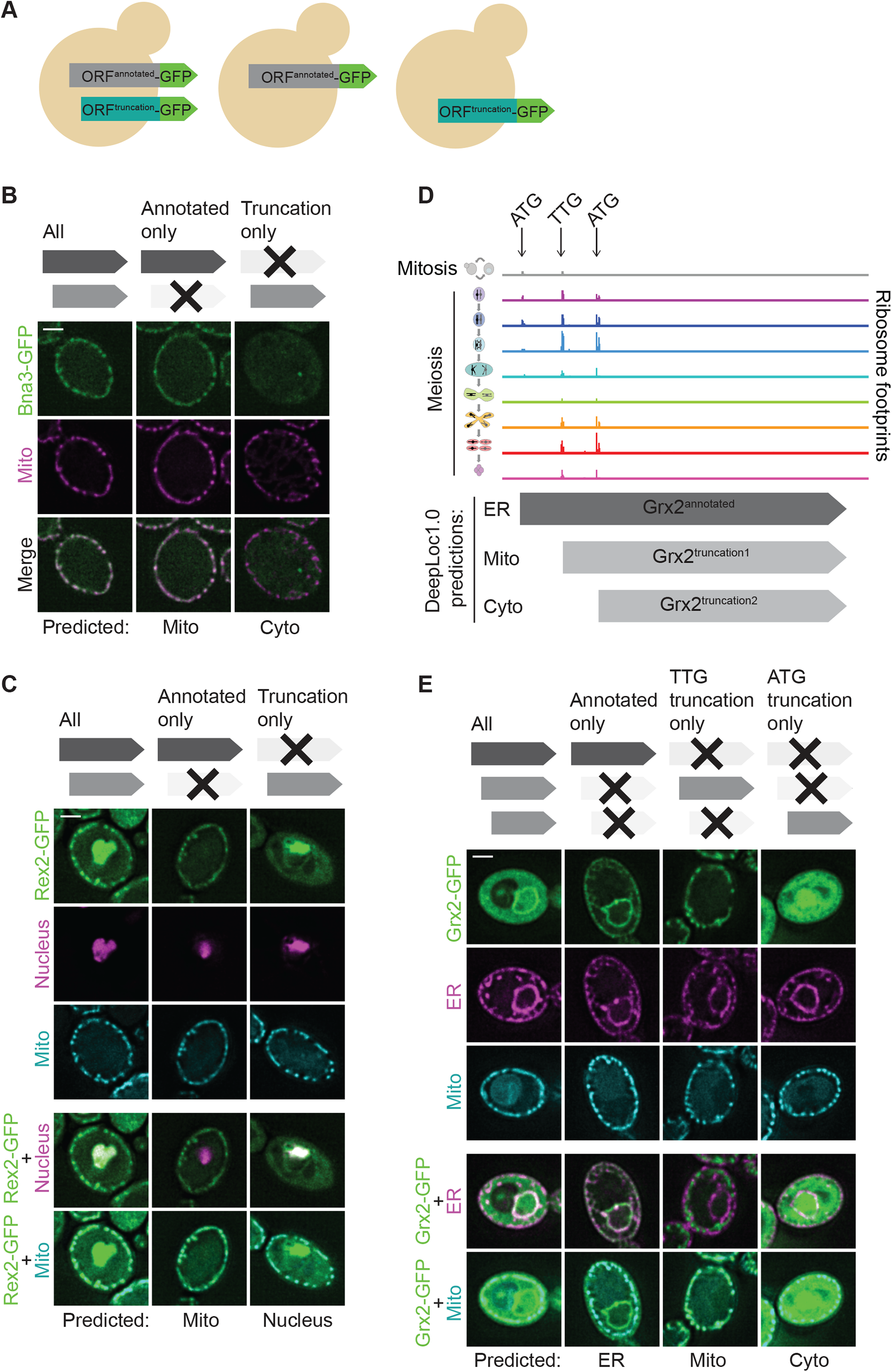
Experimental validation of differentially localized truncated isoforms. A. Schematic of experimental approach for testing localization of candidate truncated and annotated isoforms. ORFs of interest were fused to GFP to create a C-terminal tagged version of both annotated and N-terminally truncated isoforms simultaneously. Strains were constructed that contained either (1) WT start codons for both annotated and truncated isoforms, (2) the truncation start codon mutated to encode isoleucine, or (3) the annotated start codon mutated to encode isoleucine. B. Fluorescence microscopy of C-terminally GFP-tagged Bna3, collected at 6h in SPO, using the approach outlined in (A). Mitochondrial localization is indicated by Cit1-mCardinal. DeepLoc1.0-based predictions are shown below, constructs analyzed in each column are illustrated above. Scale bar is 2µm. C. Fluorescence microscopy of C-terminally GFP-tagged Rex2, collected at 1h in SPO, using the approach outlined in (A). Mitochondrial localization is indicated by Su9MTS-BFP; nuclear localization is indicated by Htb1-mCherry. Separate merged images for nuclear and mitochondrial signal are shown at the bottom. DeepLoc1.0-based predictions are shown below, constructs analyzed in each column are illustrated above. Scale bar is 2µm. D. TIS-profiling data at the *GRX2* locus across all meiotic time points. Cartoons to the left indicate whether track represents mitotic cells (gray) or meiotic timepoints (rainbow). Arrows indicate start codons for the annotated isoform and cartoons of the annotated and two predicted truncated isoforms are illustrated below, with predicted localizations for each isoform are to the left. E. Fluorescence microscopy of C-terminally GFP-tagged Grx2, collected at 0h in SPO, using an expanded version the approach outlined in (A). Strains were designed to express either all isoforms, annotated isoform only, TTG truncated isoform only, or ATG truncated isoform only. Mitochondrial localization is indicated by Su9MTS-BFP; ER localization is indicated by mCherry-HDEL. Separate merged images for ER and mitochondrial signal are shown at the bottom. DeepLoc1.0-based predictions are shown below. Constructs analyzed in each column are illustrated above. Scale bar is 2µm.

Bna3 is a kynurenine aminotransferase known to be dual localized to the mitochondria and cytoplasm but with only one characterized isoform and no known mechanism for its dual localization (Karniely et al., 2006; Wogulis et al., 2008). Since the annotated isoform was predicted to be mitochondrial and the truncated isoform was predicted to be cytoplasmic, we hypothesized that the mechanism of the previously observed dual localization could be via production of two isoforms. This prediction was indeed validated by microscopy, as the “annotated only” strain showed only mitochondrial localization and the “truncation only” strain showed cytoplasmic localization (Figure 6B).

Rex2 is an RNA exonuclease involved in snRNA, rRNA, and ncRNA processing with additional roles in mitochondrial DNA escape (Hanekamp & Thorsness, 1999; van Hoof et al., 2000). It is characterized as having mitochondrial localization, but its role in processing nuclear RNAs strongly suggested nuclear localization as well. This information aligned well with the DeepLoc1.0 prediction that the annotated isoform is mitochondrial, and the truncated isoform is nuclear. Upon imaging by microscopy, the strain containing only the annotated isoform showed mitochondrial localization and the strain containing only the truncated isoform showed nuclear localization, in support of the predicted localizations for both isoforms (Figure 6C).

Ath1 is an acid trehalase involved in extracellular trehalose degradation. Its localization has been a topic of debate, with one study showing it to be vacuolar and another showing it to be periplasmic (He et al., 2009; Huang et al., 2007). According to our predictions, the annotated isoform is Golgi-localized and the truncated isoform is extracellular. We hypothesized that the extracellular localization of the truncation could correspond to the characterized periplasmic localization that had previously been attributed to the annotated isoform. This was supported by the fact that the periplasmic localization had been observed using an endogenous C-terminal tag, which would result in tagging of both isoforms (He et al., 2009). The vacuolar localization, however, was observed with an N-terminal tag under a strong promoter, which would only tag the annotated isoform and could result in vacuolar signal simply due to protein degradation (Huang et al., 2007). Microscopy revealed that indeed the truncated isoform in isolation localizes to the periplasmic space (Figure S6E). Its levels also increase in spores, which aligns with its known function in breaking down trehalose, a storage carbohydrate that needs to be mobilized in spores. The annotated isoform, however, was not possible to visualize. This suggests that the truncated isoform is likely the main functional isoform, at least for the characterized function, and that the annotated isoform is either not produced or is at very low abundance under the conditions we assayed. This is consistent with published work showing that artificially periplasm-targeted Ath1 is sufficient for growth on trehalose while vacuole-targeted Ath1 is not (He et al., 2009).

Finally, we investigated the localization of Grx2, a glutaredoxin with two known in-frame start codons, one producing the annotated mitochondrial isoform and the second producing a truncated cytoplasmic isoform (Pedrajas et al., 2002). It was the only characterized gene with a truncated isoform for which our localization predictions did not match the characterized localizations, with the annotated isoform predicted to be ER-localized rather than the characterized mitochondrial localization (Figure 5B). To our surprise, when we imaged the annotated isoform alone, we saw clear ER localization, in line with our prediction but contradicting the established localization (Figure 5B). To reconcile this with the published data, we returned to the TIS-profiling data and noticed an additional in-frame start codon with clear signal, this time at a TTG codon, one of the most efficient of nine known near-cognate start codons (differing from ATG by one nucleotide; Figure 6D) (Kearse & Wilusz, 2017; Kolitz et al., 2008). We hypothesized that this could be the source of the missing mitochondrial localization, and indeed, the DeepLoc1.0 algorithm predicted mitochondrial localization for this isoform. To validate this prediction by microscopy, we generated strains that expressed each isoform alone by mutating the other two start codons, either to isoleucine (ATT) for the ATG starts or to a non-near-cognate leucine (TTA) for TTG. The three isoforms localized as predicted, with the annotated isoform in the ER, the TTG truncation in the mitochondria, and the ATG truncation in the cytoplasm (Figure 6E). Therefore, our truncation calling algorithm, in combination with DeepLoc1.0, correctly predicted the existence and localization of several previously characterized truncations and provided new insights into the regulation of the Grx2, a glutaredoxin that is localized to three different subcellular compartments by expression of three protein isoforms with slightly different N-termini. Together with the prediction of novel dual-localized isoforms, these localization analyses revealed that production of proximal truncations may be a widespread strategy for targeting one protein to two or more subcellular localizations.

## DISCUSSION

To date, only a handful of truncated isoforms have been characterized, and most were identified in single-gene studies of very well-studied biological pathways (Table 1). Global approaches to systematically identify these isoforms by ribosome profiling and mass spectrometry have been hampered by limitations imposed by their overlap with known proteins. Here we develop a novel algorithm for identifying truncated protein isoforms and report the translation of 388 truncated proteins in budding yeast, an organism generally considered to have little protein isoform diversity. Our analysis of this dataset represents a dramatic increase in the number of known truncated proteins in yeast and provides an unbiased picture of the prevalence and contexts in which these isoforms are present. Although functions for most remain to be investigated, we show evidence of functional activity for several truncations, with examples of proximal truncations serving to differentially localize proteins and distal truncations acting in condition-specific roles that may differ from the function of their annotated isoform. In addition to the rich global information it contains, we believe that this dataset will be a valuable resource for single gene studies, as it will provide more complete information about the gene products present at a given locus and may be helpful for generating functional hypotheses.

We orthogonally validated the production of numerous novel truncated isoforms by performing western blotting for C-terminally epitope tagged proteins (Figure 2A-G, Figure S2A-F). Of the candidates we chose, 10 were detectable by western blotting, generally at time points consistent with the TIS-profiling data. The remaining five truncations that we tested, however, were not readily detectable by western blotting despite strong and specific signal in the TIS-profiling data. We showed that several truncated proteins are subject to proteasome-mediated degradation, which in some cases made their detection in untreated cells more challenging (Figure 2K-L, Figure S2G-I). The observed degradation of this set of truncated proteins could suggest that they are turned over because they are non-functional or deleterious. It is also possible, however, that in some cases instability is an important functional feature of the protein. For one previously characterized truncation, produced at the *KAR4* locus, the annotated isoform is stable and constitutively expressed, while the truncation is much less stable and is expressed in response to mating pheromone. The unstable truncated isoform serves to provide a large boost in protein expression during mating and is then rapidly degraded when no longer needed since high levels are toxic to the cell (Gammie et al., 1999). This illustrates a scenario where the lack of stability is in fact an important feature of the truncated isoform. Such regulation could exist at other loci with truncated isoforms, providing stable and unstable pools of protein that are either functionally identical (in the case of proximal truncations) or functionally distinct (in the case of distal truncations). In general, we observed that truncated proteins beginning closer to the annotated start codon are more robustly and stably expressed than more distal truncations, which tended to be much more difficult to detect despite often having strong signal in the TIS-profiling data. We suspect that more proximal truncations are more likely to be stable since they are quite similar in size and composition to the annotated protein. Given the outsized importance of the N-terminus of proteins for protein stability, however, even a slight difference between similarly sized isoforms could confer differences in stability.

We observed dynamic regulation for most truncated isoforms in the TIS-profiling data (Figure 1F), providing support for specific cellular functions for these newly identified proteins. While TIS-profiling can give an approximate picture of regulatory patterns, it is not a robustly quantitative measure, so specific peak heights should not be overinterpreted (Eisenberg et al., 2020). We did, however, observe consistent regulatory patterns by western blotting for several examples, indicating that the TIS-profiling can be informative for regulatory trends. In many cases, this dynamic regulation is likely facilitated by the presence of a truncated transcript rather than through translational regulation. We hypothesize that truncated proteins without a detectable truncated transcript are typically produced via start codon readthrough of the annotated start codon, although this remains untested in the cases presented here (Figure 3E). It remains possible, of course, that a subset of the truncated proteins apparently lacking a truncated transcript in fact arise from false positives in the TIS-profiling or false negatives in the TL-seq data; however, we expect that this would be a minor contribution.

A large subset of the proximal truncations that we identified seem to lead to otherwise functionally identical proteins being targeted to different subcellular locations, which provides a mechanism for how in so many cases, similar cellular functions are performed in multiple cellular compartments. For example, DNA replication, transcription, and translation all occur in both the mitochondria and either the nucleus or cytoplasm. These types of related but spatially separated functions can be encoded either by two separate genes (through gene duplication or functional convergence) or by a single gene (Danpure, 1995). Previous single-gene studies made it clear that a single locus can encode multiple differentially localized protein isoforms, but the full extent of the phenomenon was unknown, and the known examples were biased towards very well-studied biological pathways (Table 1). For example, several previously characterized truncated isoforms were amino acid tRNA synthetases, known to act on both cytoplasmic and mitochondrial tRNAs, and were therefore clear candidates for this type of regulation. Here, genome-wide data allowed us to see the extent of this type of regulation in yeast in a less biased way. Candidates that we validated for their predicted localization differences spanned a diverse range of functions and revealed a variety of ways in which knowledge of this cellular strategy can enhance our understanding of gene function and regulation.

In the case of Bna3, dual protein localization was already established, but its basis was unknown. The two isoforms at this locus follow a common trend seen among previously characterized truncations, in which the longer form has a mitochondrial signal sequence that is lost in the truncated isoform, causing it to default to cytoplasmic localization (Figure 6B). *REX2* presented a slightly different scenario, in which dual localization had not been established but was very likely given known functional information. Rex2 had been characterized as a mitochondrial protein by mitochondrial fractionation of overexpression strains (Hanekamp & Thorsness, 1999). Despite not being explicitly characterized as nuclear by microscopy, it was also shown to be involved in rRNA and snRNA processing, strongly suggesting a nuclear localization (van Hoof et al., 2000). Our predictions and validation support both localizations and suggest that the mechanism of dual localization is through production of two differentially targeted protein isoforms (Figure 6C). It is also an interesting example of one signal sequence being removed (mitochondrial) and another being unmasked (nuclear) in the truncated isoform. Interestingly, we did not find dual localization for *ATH1* and instead showed that the characterized function is likely carried out by the truncated isoform, and that the annotated isoform is likely not expressed at appreciable levels (Figure S6E). Identification of the truncated isoform and subsequent localization predictions were valuable for explaining inconsistencies in the existing literature.

The *GRX2* locus provides an additional demonstration of how TIS-profiling can help reconcile confusing information about a gene’s regulation and function. Based on previous characterization, *GRX2* was thought to produce two protein isoforms, one mitochondrial and one cytoplasmic (Porras et al., 2006). In that study, however, three bands were observed by western blotting, which likely correspond to the three isoforms that we identified. The long isoform, however, was hypothesized to be mitochondrial and the intermediate isoform was attributed to processing of the mitochondrial targeting sequence from the longer isoform. A small amount of protein was also detected in the ER, but this was attributed to slow import kinetics into the mitochondria. With the additional insight provided by the TIS-profiling paired with localization prediction, we were able to identify a third isoform of the redox-regulator Grx2. Visualization of the protein structure using AlphaFold shows that all three isoforms, which localize to three different cellular compartments, still retain the structured, functional core of the protein (Figure S6F, Figure 6D-E) (Jumper et al., 2021).

The identification of a TTG-initiated truncation at the *GRX2* locus raises the question of whether near-cognate start codons should have been included in our truncation calling algorithm. We chose to exclude them because visual analysis indicated that most near-cognate truncations were false positives caused by background noise within genes and would likely require different treatment and calling thresholds than AUG start sites. While the *GRX2* locus shows that near-cognate-initiated truncations can be made, we still believe that they are very rare. Translation at near-cognate start codons alone is often not sufficient to stabilize a transcript, as shown in past work in which mutation of the annotated start codon of transcripts encoding a near-cognate-initiated extended protein isoform led to nonsense-mediated decay of the transcript; this was caused by efficient translation initiation at an out-of-frame AUG codon downstream following inefficient initiation at the in-frame near-cognate codon (Eisenberg et al., 2020). Therefore, we suspect that most cases of near-cognate truncations would need to arise in a context similar to *GRX2*, in which a truncated transcript bears an ATG truncation, whose translation ensures the stability of the transcript, paired with an upstream near-cognate-initiated isoform. In these cases, the near-cognate isoform is essentially behaving like an N-terminal extension within the context of the truncated transcript.

We observed dynamic and condition-specific regulation for two distal truncations that we investigated experimentally, Yap5^truncation^ and Pus1^truncation^, which suggests functional relevance. Yap5^truncation^ contains the Fe-S cluster binding domain located in the C-terminal half of the annotated protein and is markedly similar in size to an artificial truncation of Yap5 that was shown to effectively bind Fe-S clusters (Rietzschel et al., 2015). We show that Yap5^truncation^ is specifically induced under multiple respiratory conditions: meiosis, saturated growth, and growth in non-fermentable media (Figure 2A, Figure 4A-B). This suggests that it may be involved in responding to elevated respiratory activity, a role which could be related to its ability to bind Fe-S clusters, important cofactors in the electron transport chain. Further work will be necessary to elucidate its specific functional role.

Pus1^truncation^ is produced throughout meiosis and contains the positively charged residues involved in RNA binding of the full length protein (Czudnochowski et al., 2013). We show that this truncated isoform is likely nutrient-regulated since it is expressed in the low nutrient media that induces meiosis, as well as upon glucose starvation and rapamycin treatment (Figure 2D, Figure 4D-E). This nutrient regulation is intriguing given that a number of Pus1-dependent modifications in mRNA are dynamically regulated during nutrient deprivation (Carlile et al., 2014; Schwartz et al., 2014). The induction of Pus1^trunction^ upon rapamycin treatment is also interesting; formation of some pseudouridines by other Pus proteins is known to be dynamically regulated by the TOR pathway, and a pseudouridine in the U6 snRNA is introduced by Pus1 during filamentous growth, also regulated by the TOR pathway (Basak & Query, 2014; G. Wu et al., 2011; G. Wu, Radwan, et al., 2016).

mRNA-seq of cells expressing Pus1^truncation^ in WT and *pus1Δ* backgrounds under rich growth conditions revealed mild effects of either Pus1^truncation^ expression or *PUS1* deletion alone, and a much more dramatic effect when deletion of *PUS1* is combined with Pus1^truncation^ expression (Figure 4F-G). This more severe synthetic gene expression phenotype suggests that Pus1^truncation^ has effects that are not directly related to full-length Pus1 function, potentially also affecting targets of other pseudouridine synthases as well. This is perhaps unsurprising given that the specificity of the enzyme is primarily conferred by the catalytic domain and there is little reason to think the RNA binding domain alone would be specific to Pus1 targets.

The precise reason for the strong downregulation of genes involved in ribosome biogenesis, rRNA processing, and ncRNA processing in cells expressing Pus1^truncation^ and deleted for *PUS1* is not immediately obvious (Figure S4D). Pseudouridine synthases as a group perform extensive pseudouridylation of rRNA, tRNA, snRNA, snoRNA, and mRNA targets, any of which could have important impacts on translation and RNA processing (reviewed in Rintala-Dempsey & Kothe, 2017). Pus1 itself has been shown to modify ribosomal mRNAs, including 5 subunits of the ribosomal large subunit, as well as RNase MRP which is involved in maturation of rRNA (Carlile et al., 2014; Schwartz et al., 2014). Given that Pus1^truncation^ contains regions involved in RNA-binding, we hypothesize that it could occlude target binding by Pus1 and potentially other pseudouridine synthases as well (Czudnochowski et al., 2013). The presence of a synthetic effect with deletion of *PUS1* is perhaps suggestive of a role in modulating pseudouridylation. While deletion of *PUS1* alone results in viable cells and only mild phenotypic affects, much more dramatic synthetic phenotypes have been observed when *PUS1* deletion is combined with deletion of other pseudouridine synthases or with mutations that compromise tRNA stability (Großhans et al., 2001; Khonsari & Klassen, 2020; G. Wu, Adachi, et al., 2016). If Pus1^truncation^ has a role related to pseudouridylation, a synthetic phenotype with deletion of *PUS1* would be consistent. Further study will be necessary to understand the specific mechanistic role of _Pus1_truncation.

Our hypotheses for Yap5^truncation^ and Pus1^truncation^ function were notably tied to the known functional characteristics of their annotated isoforms. Whether this is a valid approach is unclear, as many truncations – particularly distal truncations – lack key functional sequences of the annotated protein. The degree to which the annotated and truncated isoform differ in function likely varies depending on the extent of the truncation, with the proximal truncations being much more likely to share functional characteristics with the annotated isoform and only varying in typical N-terminally encoded characteristics such as stability and localization. Distal truncations, on the other hand, are more likely to be missing key functional domains and may not even contain any intact domains, making it much more difficult to generate a rational prediction for their functions.

Beyond the new regulation we uncovered for Pus1 as a result of identifying its truncated isoform, this case highlights a key point: our truncation-calling algorithm is stringent, likely excluding a number of real truncated isoforms in order to minimize false-positive calls, which previously plagued identification of N-terminal truncations. Systematic identification of this type of non-canonical protein is fundamentally distinct from other classes, including ORFs in upstream regions (uORFs), those that are short and intergenic (sORFs), those downstream of annotated ORFs (dORFs), and N-terminally extended ORFs. In all other cases, some or all of the novel ORF is non-overlapping with an annotated ORF. Thus, approaches based on standard ribosome profiling data that leverage initiation codon peaks resulting from cycloheximide pre-treatment, periodicity resulting from elongation, or simply ribosome density, are not effective for stringently calling N-terminal truncations. Even approaches independent of standard ribosome profiling, like those that use evolutionary conservation to identify coding regions, are problematic for this class in particular; and those that rely on machine learning-based analysis of translation initiation site mapping in conjunction with standard ribosome profiling analysis generate an exceedingly high level of false positives and negatives, likely due to multiple of the factors noted above (Eisenberg et al., 2020). Our systematic identification of truncated proteins, in contrast, relied entirely on TIS-profiling data. Relative to standard ribosome profiling, the TIS-profiling data is much simpler to interpret since the reads are highly enriched at sites of translation initiation and signal is not obscured by elongating ribosomes from the overlapping annotated ORF, allowing us to very robustly identify truncated protein initiation sites (Figure 1A-B). Analysis of parallel standard ribosome profiling data provided clear validation of our calls, suggesting that future studies can rely on TIS-profiling for protein isoform identification (Figure 1E). Furthermore, multiple lines of experimental testing revealed that these isoforms can have localization and function distinct from their corresponding annotated ORF.

Non-canonical translation has previously been shown to be higher during meiosis and other stress conditions, and from our observations in this dataset, truncated isoforms are no exception (Brar et al., 2012; Cheng et al., 2018; Eisenberg et al., 2020). This could be evolutionarily beneficial, allowing cells to sample a greater proteomic diversity to adapt to new environments. Some truncated isoforms may not currently be “useful” to cells but may eventually over time become functional. While the use of different isoforms bears some similarity to gene duplication, it is markedly different in that the shared sequences between the two isoforms are unable to evolve separately. Only the region missing in the truncation can change independently between the two isoforms. N-terminal sequences, however, are often particularly important for gene function, as we have shown for localization and stability. Therefore, having a mechanism to test out different N-terminal sequences while still retaining protein production from the annotated start site could be beneficial. Future work on the prevalence and conservation of truncated isoforms across different stress conditions and other organisms will further elucidate both the functional relevance and evolutionary processes giving rise to truncated proteins.

## FIGURE LEGENDS

**Figure S1.**
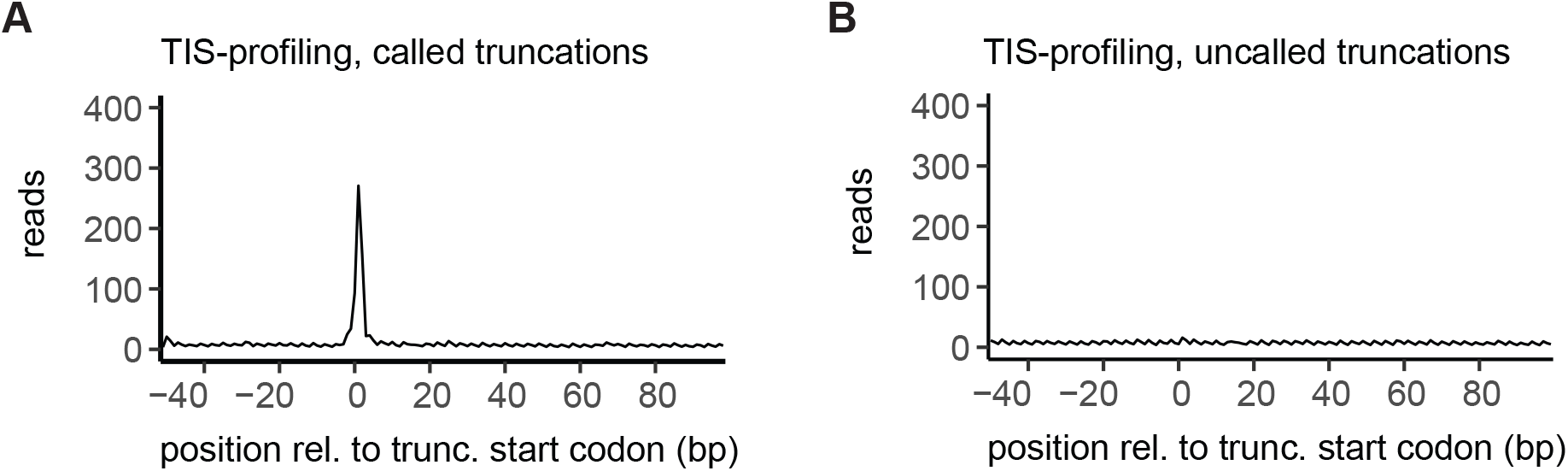
A. Metagene plot of TIS-profiling data for all called truncated isoforms for the region between -50 and +100bp relative to the truncation start codon. Reads are summed across all timepoints. B. Metagene plot of TIS-profiling data for all uncalled truncated isoforms for the region between -50 to +100bp relative to the truncation start codon. Reads are summed across all timepoints.

**Figure S2.**
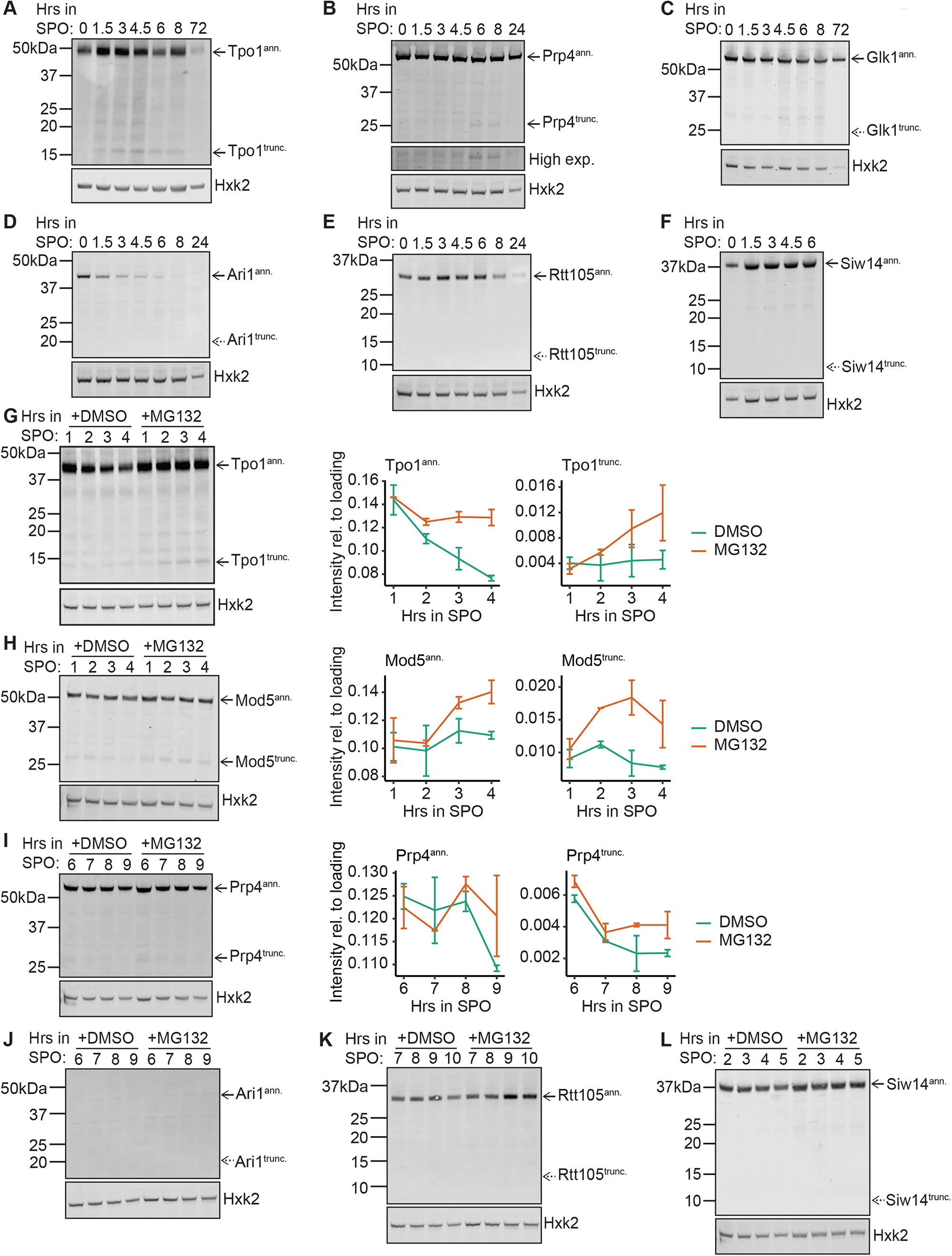
A. Western blot of samples collected at various timepoints after transfer of cells to sporulation media (SPO). Hexokinase (Hkx2) is shown as a loading control. A high exposure (high exp.) panel is included for lowly expressed truncations. A different strain is used in each case, expressing the indicated C-terminal epitope-tagged protein and enabling detection of annotated and truncated isoforms for (A) Tpo1-3V5, B. (B) Prp4-3V5, C. (C) Glk1-3V5, D. (D) Ari1-3V5, E. (E) Rtt105-3V5, F. (F) Siw14-3V5, G. Representative western blot (left) and quantification (right) for cells treated with proteasome inhibitor MG132 or vehicle control DMSO. Quantification is based on 2 replicates and error bars represent standard error. Hexokinase (Hkx2) is shown as a loading control. Blots show stabilization of truncated isoforms for (G) Tpo1-3V5, H. (H) Mod5-3V5, I. (I) Prp4-3V5, J. Western blot analysis for cells treated with proteasome inhibitor MG132 or vehicle control DMSO. Hexokinase (Hkx2) is shown as a loading control. Blots show lack of stabilization of truncated isoforms for (J) Ari1-3V5, K. (K) Rtt105-3V5, L. (L) Siw14-3V5

**Figure S3.**
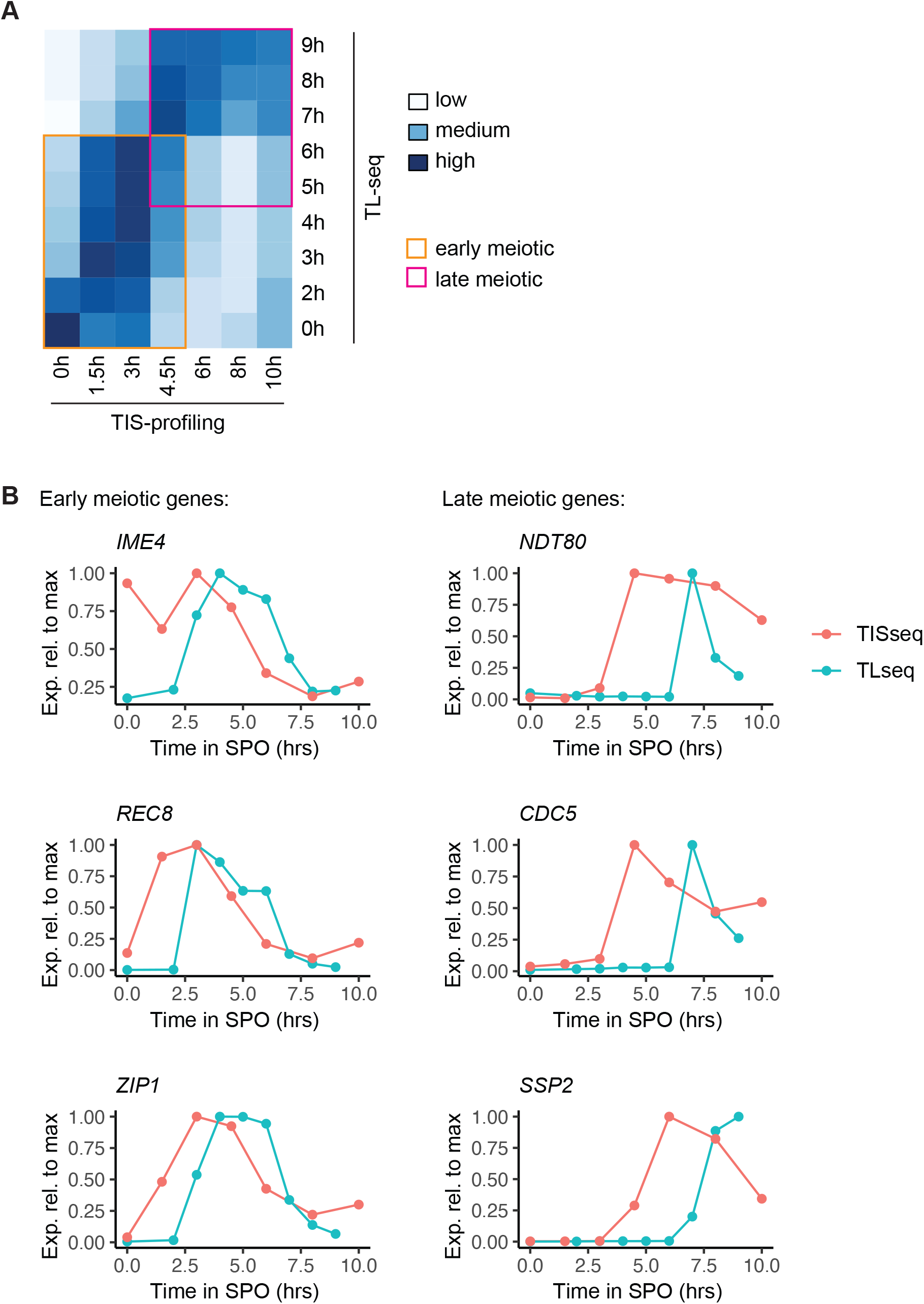
A. Heatmap of Pearson correlations between the two mRNA-seq time courses collected in parallel with the TL-seq and TIS-profiling data, used to compare meiotic staging between the two time courses. Meiotic time points are labeled along the x and y axes. Early and late meiotic time point groups are boxed in orange and pink, respectively. B. Plots of the expression relative to max for example early (left) and late (right) meiotic genes from mRNA-seq time courses collected in parallel with the TL-seq and TIS-profiling data.

**Figure S4.**
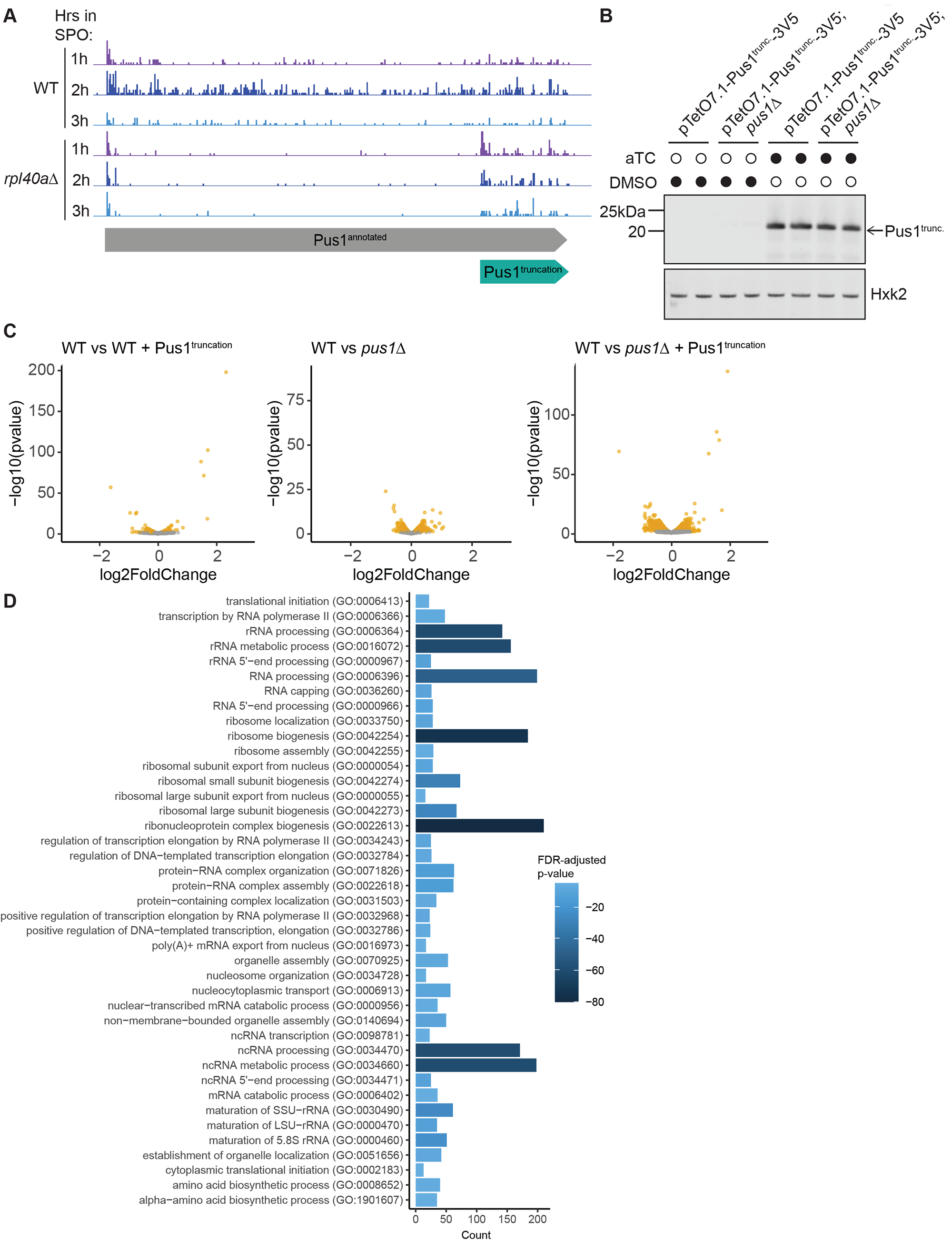
A. Standard ribosome profiling data of WT and *rpl40aΔ* cells at the *PUS1* locus. Cartoons below represent the annotated and truncated isoforms of Pus1. Samples were collected at indicated timepoints following transfer to sporulation media (SPO). B. Western blot confirming expression of Pus1^truncation^ upon aTC treatment. Samples were collected in WT and *pus1Δ* cells carrying a construct to allow anhydrotetracycline-inducible Pus1^trunc.^ expression, treated with either vehicle (DMSO) or aTC. C. Volcano plot of DESeq2 analysis of mRNA-seq data for the following conditions: WT cells carrying a construct to allow aTC-inducible Pus1^trunc.^ expression and treated with aTC (“WT + Pus1^trunc.^”), *pus1Δ* cells carrying a construct to allow aTC-inducible Pus1^trunc.^ expression and treated with vehicle (“*pus1Δ”*) or aTC (“*pus1Δ* + Pus1^trunc.^*”*). In all cases differential expression is relative to WT cells carrying a construct to allow aTC-inducible Pus1^trunc.^ expression and treated with vehicle (“WT”). Points for significantly differentially expressed genes, as called by DESeq2 (p-adj < 0.1), are yellow and non-significant genes are gray. D. Top hits from GO term analysis of significantly downregulated genes between WT and *pus1*Δ + Pus1^truncation^ cells.

**Figure S5.**
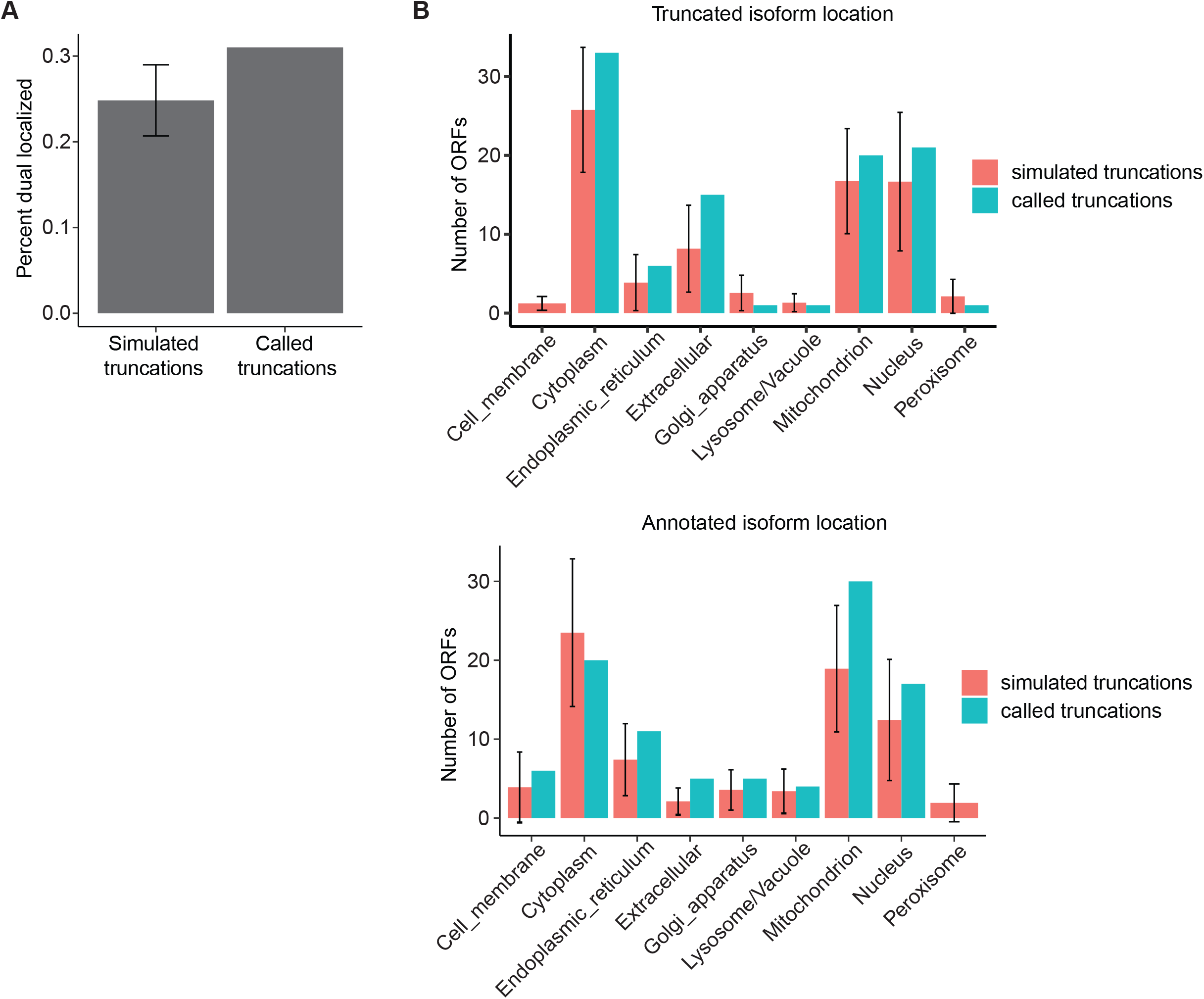
A. Bar plot of percent of truncated isoforms that are differentially localized relative to their annotated isoform, compared to the percent of simulated truncations (randomly sampled in-frame start codons). Error bar represents 2 standard deviations. B. Bar plot of number of truncated (upper) or annotated isoforms (lower) localized to each subcellular compartment, compared to simulated truncations (randomly sampled in-frame start codons). Error bars represent 2 standard deviations.

**Figure S6.**
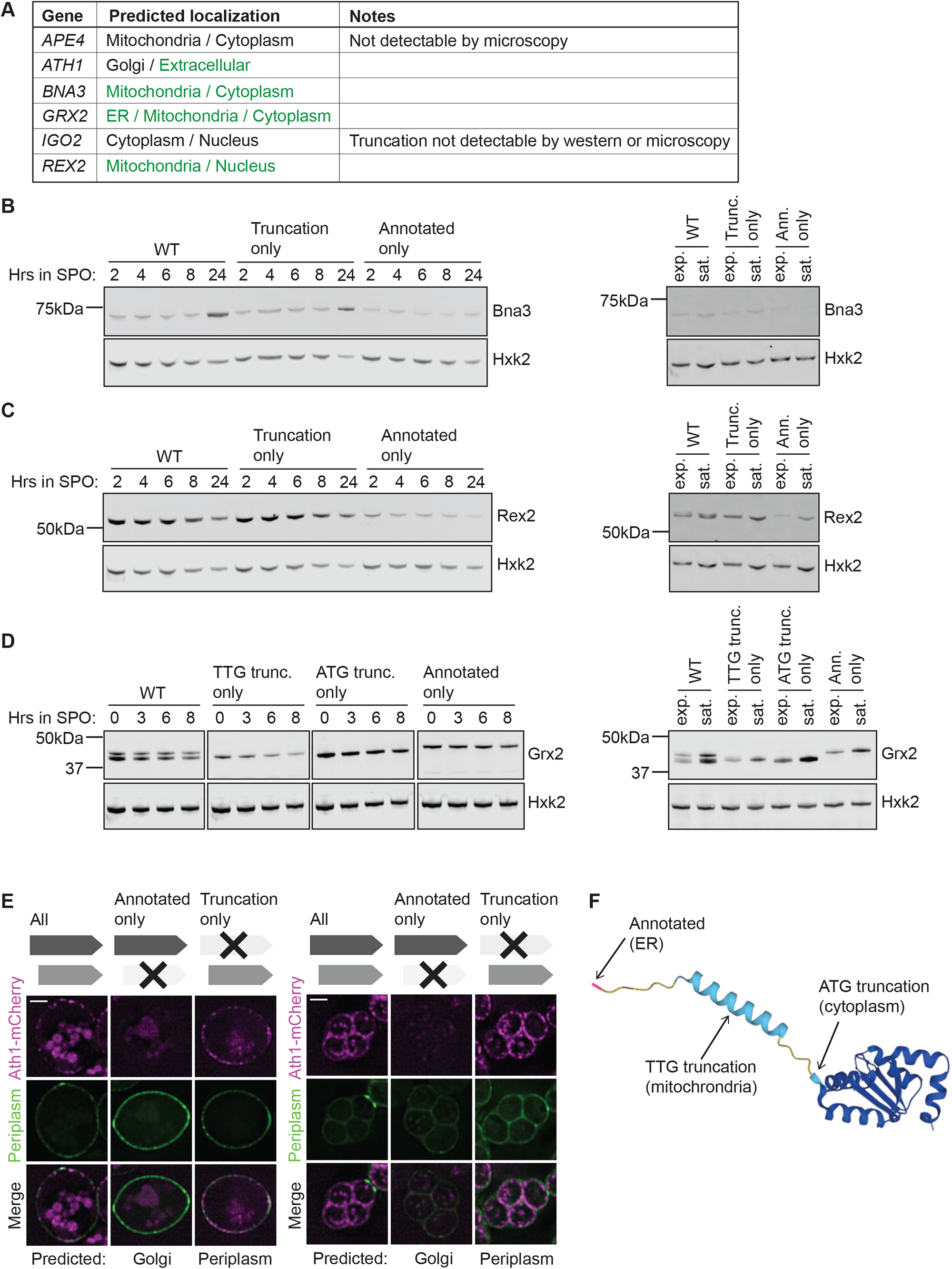
A. Table of candidates chosen for validation by microscopy, including their predicted localizations (annotated / truncation). Green represents localization predictions that were successfully validated, black represents unvalidated predictions. B. Western blot of samples collected at various timepoints after transfer of cells to sporulation media (SPO) or in vegetative exponential or vegetative saturated growth for strains described in Figure 6A. Hexokinase (Hxk2) is shown as a loading control. Blots are for (B) Bna3-GFP, C. (C) Rex2-GFP, D. (D) Grx2-GFP, for strains described in Figure 6E E. Fluorescence microscopy of C-terminally mCherry-tagged Ath1, collected at 3h (left) and 24h (right) in SPO, using the approach outlined in (6A). Periplasmic localization is indicated by Suc2-GFP. DeepLoc1.0-based predictions are shown below, constructs analyzed in each column are schematized above. Scale bar is 2µm. F. Alpha fold structural prediction for full-length Grx2, with arrows indicating the residues corresponding to the start codons of the annotated (ER), TTG truncation (mitochondrial), and ATG truncation (cytoplasmic) isoforms.

## SUPPLEMENTARY FILES

Table S1. List of called truncated protein isoforms.

Table S2. List of truncated isoforms called as having a TSS based on TL-seq data.

Table S3. DeepLoc1.0 localization predictions for truncated isoforms and their annotated counterpart.

Table S4. Pus1 RNA-seq data, including the raw counts matrix, RPKMs, and DEseq output. Table S5. Gene ontology analysis for clusters of differentially expressed genes in Pus1 dataset. Table S6. List of yeast strains and plasmids used in this study.

## MATERIALS AND METHODS

### Yeast strain construction

Strains were constructed in the SK1 background of *Saccharomyces cerevisiae*. Strains and plasmids used for this study are listed in Table S6. Deletion strains were created using pÜB81, and C-terminal 3V5 or FLAG-tagged strains were generated via Pringle tagging at the endogenous locus using pÜB81 or pÜB166 (Longtine et al., 1998). GFP-tagged strains for microscopy were generated using PmeI-digested single-integration plasmids constructed via Gibson assembly of PCR-amplified fragments containing the ORF of interest along with its own 5’ leader region amplified from genomic DNA and backbone fragments containing either a GFP tag, *ADH1* terminator, and a *TRP1* selection marker (pÜB629) or an mCherry tag, *ADH1* terminator, and a *HIS3* selection marker (pÜB1736). Start codon mutants were generated from single-integration plasmids described above by PCR amplifying fragments for Gibson assembly using primers containing the desired point mutation. Pus1^truncation^ overexpression strains were generated following the WTC_846_ system (Azizoglu et al., 2021). The truncated open reading frame sequence was inserted into a single-integration plasmid downstream of pTetO7.1 by Gibson assembly of PCR fragments from genomic DNA and backbone fragments from pUB2344. Transformants were crossed into strains containing pRNR2-TetR-Tup1 and pTetO7.1-TetR.

### Yeast growth and sporulation

For vegetative experiments, strains were grown in YEPD at 30°C. Strains were inoculated and grown overnight to reach saturation (OD600 > 10), then back-diluted to an OD600 of 0.2 and grown to desired OD600. For meiotic time courses, strains were inoculated into YEPD supplemented with uracil and tryptophan (1% yeast extract, 2% peptone, 2% glucose, 22.4 mg/L uracil, and 80 mg/L tryptophan) and grown for 24h at RT to an OD600 ≥ 10, then diluted to an OD600 of 0.25 in buffered YTA (1% yeast extract, 2% bacto tryptone, 1% potassium acetate, and 50 mM potassium phthalate) and grown for 16h at 30°C to an OD_600_ ≥ 5. Cells were spun down and washed once with sterile MilliQ water before resuspension in sporulation media (SPO; 2% potassium acetate supplemented with amino acids (40 mg/L adenine, 40 mg/L uracil, 10 mg/L histidine, 10 mg/L leucine and 10 mg/L tryptophan)) at OD600 = 1.85 and shaken at 30°C, with timepoints collected at times indicated in figures.

### Protein extraction and western blotting

Strains were grown in specified media and 2 or 3.3 OD600 equivalents of cells were collected for vegetative and meiotic cultures, respectively. Samples were incubated in 5% TCA for ≥10mins at 4*°*C then spun down, washed once with TE, once with acetone, then dried overnight. Pellets were resuspended in 150ul of lysis buffer (50mM Tris-HCl, 1mM EDTA, 3mM DTT, 1.1mM PMSF (Sigma), and 1X cOmplete mini EDTA-free protease inhibitor cocktail (Roche)) and cells were lysed by bead-beating for 5min at RT. SDS loading buffer was added to 1X and samples were incubated at 50°C for 10min and beads were pelleted by centrifugation. Samples were run on a 4-12% Bis-Tris gel at 160V for 5min followed by 200V for 25min. Transfer to nitrocellulose membrane was performed using a semi-dry transfer system (Trans-Blot Turbo, BioRad) with a standard 30 min transfer. The membrane was blocked in 5% milk PBS-T for 1 hour at RT and incubated in primary antibody overnight at 4°C. Primary antibodies were diluted in 5% milk in PBS-T + 0.01% sodium azide (1:2,000 for mouse anti-GFP (Clontech) and mouse anti-3V5 (Invitrogen), 1:1000 for mouse anti-FLAG (Sigma), and 1:10,000 for rabbit anti-hexokinase (Rockland). Membrane was washed 3X in PBS-T then incubated in secondary antibody (1:15,000 anti-mouse 800 and anti-rabbit 680 in LI-COR PBS blocking buffer) for 1 hour at RT, then washed 3X in PBS-T before imaging on the LI-COR Odyssey Imager. Analysis and quantification was performed using ImageStudio Lite software.

### Proteasome inhibition

Strains were constructed in a *pdr5Δ* background to confer drug sensitivity. Standard meiosis conditions were used as described above. At the designated time point, cultures were split into two cultures and 100uM MG132 or DMSO (vehicle) was added. Cells were collected for protein extraction and western blot as described above at time points indicated in figures.

### Growth in non-fermentable media

Cells were grown to saturation overnight in YEPD then back-diluted to an OD600 of 0.2 in YEPD. At 4h post-dilution, cells were spun down and resuspended in either YEPD (fermentable) or YEPG (non-fermentable). Cells were collected for protein extraction and western blot as described above at time points indicated in figures.

### Rapamycin treatment

Cells were grown to saturation overnight in YEPD then back-diluted to an OD600 of 0.2 in YEPD. At 2 hours, cultures were split and treated with either rapamycin (0.2ug/ml or 0.5 ug/ml) or DMSO (vehicle). Cells were collected for protein extraction and western blot as described above at time points indicated in figures.

### Low glucose growth

Cells were grown to saturation overnight in YEPD then back-diluted to an OD600 of 0.2 in either YEPD with 2% dextrose (normal) or 0.2% dextrose (low). Cells were collected for protein extraction and western blot as described above at time points indicated in figures.

### Pus1^truncation^ overexpression

Cultures were grown to saturation overnight in YEPD then back-diluted to an OD600 of 0.2 in YEPD and treated immediately with either 1ug/ml anhydrotetracycline (aTC) or DMSO (vehicle). Samples were collected 3h post-dilution.

### RNA extraction

5ODs of cells were pelleted by centrifugation and flash frozen in liquid nitrogen. Cells were thawed on ice and resuspended in TES buffer (10 mM Tris pH 7.5, 10 mM EDTA, 0.5% SDS). An equal volume of acid phenol (pH4.3, Sigma-Aldrich) was added. Samples were shaken at 1400rpm for 30min at 65°C, then spun down at 4°C. The aqueous phase was transferred to a new tube containing 350ul chloroform. Samples were spun down and the aqueous layer was transferred to a new tube containing 100% isopropanol and with 350mM sodium acetate (pH5.2). Samples were precipitated overnight at -20°C. RNA was pelleted by centrifugation and pellets were washed with 80% ethanol, dried, resuspended in DEPC water for 10min at 37°C. Total RNA was quantified using the Qubit RNA BR Assay Kit (ThermoFisher).

### Poly-A selection and RNA-seq

Poly-A selection was performed using the NEXTFLEX Poly(A) Beads 2.0 kit with 5ug total RNA (NOVA-512992). RNA-seq libraries were prepared from the resulting poly-A selected RNA using the NEXTFLEX Rapid Directional RNA-Seq Kit 2.0 (NOVA-5198-02). Libraries were quantified and quality checked using the Agilent 4200 TapeStation (Agilent Biotechnologies Inc). Samples were sequenced on the NovaSeqX sequencer.

### Live imaging

At designated time points 2ul of meiotic culture was placed on a glass slide and imaged immediately. Images were acquired using a DeltaVision Elite wide-field fluorescence microscope (GE Healthcare), a 100X/1.40 NA oil-immersion objective (DeltaVision, GE Healthcare, Sunnyvale, CA), and the following filters: FITC, mCherry, DAPI. 30 z-stacks were collected with 0.2uM spacing. Images were deconvolved using softWoRx imaging software (GE Healthcare).

### Sequence alignment, quantification, and differential expression analysis

Sequencing data were aligned to the SK1 genome using STAR. A-site mapping for standard ribosome profiling and TIS-profiling data was performed as previously described (Eisenberg et al., 2020). Differential expression analysis for mRNA-seq data was performed using DESeq2. Hierarchical clustering was performed using complete-linkage clustering on the Pearson correlation of the log2-transformed average of 2 replicates. Genome browser visualization was performed using IGV.

### Gene ontology enrichment analysis

GO analysis was performed using the PANTHER classification system (Mi et al., 2013).

### Truncation calling algorithm

Analysis was performed on TIS-profiling data collected at 0h, 1.5h, 3h, 4.5h, 6h, 8h, 10h, and 22h after addition to sporulation media (SPO), as well as in vegetative exponential, vegetative saturated growth and a MATa/a non-meiotic starvation control collected at 4.5h in SPO, as described in (Eisenberg et al., 2020). To generate a list of putative truncation-generating start codons, we first found all in-frame start codons within annotated exons. For each potential start codon (ATG) at each timepoint, a “peak sum” was calculated by summing the reads at the three nucleotides corresponding to the start codon. To model the background reads for each gene at each time point, we generated an empirical distribution of peaks sums from sets of three independent nucleotides that were randomly sampled with replacement (10,000x). The empirical p-value for each putative start codon, including annotated start codons, was determined by comparing the peak sum for the codon of interest to the empirical distribution. Annotated and putative truncation start codons were then filtered with the following criteria: p-value ≤0.0015 and >11 reads for at least one nucleotide in the start codon. To be considered in the final set, each truncation was required to be called at 2 or more timepoints. Putative truncations were additionally required to start ≥5aa from the annotated start codon and have an ORF length >10aa. Cases of likely mis-annotation, where the “truncated” isoform is likely the dominant isoform, were also removed; this gene set was generating through computational filtering to identify genes where the annotated isoform was not called followed by manual curation through visualization in a genome browser.

### Ribosome profiling metagene analysis

Reads were averaged across all truncations at all positions between -50bp and +100bp surrounding truncated isoform TISs. We excluded truncations that begin within 50bp of the annotated isoform to avoid including reads associated with annotated start peaks that would confound the 5’ signal. To prevent the profile from being overpowered by single highly expressed genes, we excluded genes with a Z-score greater than 10 at any position.

### TL-seq metagene analysis

Reads were summed across all truncations at all positions between -200bp and +200bp surrounding annotated or truncated isoform TISs. We excluded truncations that begin within 200bp of the annotated isoform to avoid including reads associated with annotated TSSs. To prevent the profile from being overpowered by single highly expressed genes, we excluded genes with a Z-score greater than 10 at any position.

### TL-seq peak calling

Counts per site were extracted from published bigwig files using custom scripts (Chia et al., 2021). To call protein isoforms with 5’ transcript ends upstream of their start codons, for each gene an upstream-to-downstream ratio was calculated, such that { ratio = sum(reads 200bp upstream)/sum(reads 200bp downstream) }. Each gene’s upstream-to-downstream ratio was compared to an empirical distribution of 10,000 random ratios obtained by taking the ratio of the sums of two randomly sampled groups of 200 sites within the gene, sampled with replacement. Reads upstream of the annotated TIS were masked to avoid including reads derived from 5’ ends of annotated transcript isoforms. A p-value was calculated using the empirical cumulative distribution function of these ratios, and a p-value cutoff of 0.1 was used. To exclude genes with very sparse or no coverage we additionally required a variance greater than 0.05 for the distribution of sample ratios.

### Staging comparison between TIS-profiling and TL-seq time courses

Stage matching between time courses was performed using an mRNA-seq time course collected in parallel with the TL-seq time course and an mRNA-seq time course collected under matched strain and growth conditions as the TIS-profiling time course. Note that staging was performed differently for the two time courses – the TIS-profiling was synchronized naturally via starvation conditions, whereas the TL-seq time course was synchronized via inducible expression of meiotic master regulator transcription factors *IME1* and *NDT80*. See (Cheng et al., 2018; Chia et al., 2021) for details. Timepoints for each time course were split into either early-meiotic (TIS-profiling: 0h, 1.5h, 3h, 4.5h; TL-seq: 0h, 2h, 3h, 4h, 5h, 6h) or late-meiotic (TIS-profiling: 4.5h, 6h, 8h, 10h; TL-seq: 5h, 6h, 7h, 8h, 9h) based on Pearson correlation of log2-transformed RPKMs and expression patterns of key meiotic genes in the mRNA-seq time courses (Figure S3A-B).

### Localization prediction

DeepLoc1.0 was run on the amino acid sequence of all called truncated isoforms as well as their corresponding annotated isoform (Almagro Armenteros et al., 2017). As a control, we generated sets of simulated truncations by randomly sampling in-frame ATGs within annotated genes that are not called as real start sites. To ensure that the length distribution of the control set approximately matched the set of real truncations, for each real truncation we randomly sampled an in-frame start with the distance from annotated isoform within +/-5 amino acids of that of the real truncation.

### Resource availability

All reagents used in this study are available upon request from the corresponding author. Sequencing data will be made available at NCBI GEO. Custom analysis code will be made available on GitHub.

## ACKNOWLEDGEMENTS

We thank Nick Ingolia and members of the Brar and Ünal labs for critical feedback on the manuscript. We also thank Jeremy Thorner, Amy Eisenberg, Emily Powers, and Josh Schraiber for helpful conversations and insights about the project. This work was supported by National Institutes of Health [1R35GM134886] funding to GAB. ALH was also funded by an NSF predoctoral fellowship [DGE 1752814] and an NIH training grant to the MCB department [GM T32 007232].

## AUTHOR CONTRIBUTIONS

ALH and GAB conceived most aspects of this study. Western blot validation of truncations and proteasome inhibition experiments were performed by ALH and NW. All other experiments were performed by ALH. Algorithm development and data analysis was performed by ALH. The manuscript was written and edited by ALH and GAB.

## DECLARATION OF INTERESTS

The authors declare no competing interests.

## Notes

### Competing Interest Statement

The authors have declared no competing interest.

